# Structural and Thermodynamic Impact of Oncogenic Mutations on the Nucleosome Core Particle

**DOI:** 10.1101/2025.02.14.638149

**Authors:** Augustine C. Onyema, Christopher DiForte, Rutika Patel, Sébastien F. Poget, Sharon M. Loverde

## Abstract

The nucleosome core particle (NCP) is essential for chromatin structure and function, serving as the fundamental unit of eukaryotic chromatin. Oncogenic mutations in core histones disrupt chromatin dynamics, altering DNA repair and transcription processes. Here, we investigate the molecular consequences of two mutations—H2BE76K and H4R92T—using 36 µs of all-atom molecular dynamics simulations and experimental biophysical assays. These mutations destabilize the H2B-H4 interface by disrupting critical salt bridges and hydrogen bonds, reducing binding free energy at this interface. Principal component analysis reveals altered helix conformations and increased interhelical distances in mutant systems. Thermal stability assays (TSA) and differential scanning calorimetry (DSC) confirm that these mutations lower the dimer dissociation temperature and reduce enthalpy compared to the wild type. Taken together, our results elucidate how these mutations compromise nucleosome stability and propose mechanisms through which they could modulate chromatin accessibility and gene dysregulation in cancer.

**Statement of Significance:** The nucleosome is the essential packaging unit of DNA. ‘Oncohistones’ which are histone mutations that are associated with cancer, are known to compromise the stability of the nucleosome and affect nucleosome sliding. Here we perform long-time molecular dynamics simulations of two buried histone core mutations, demonstrating that these mutations reduce the stability of the histone core at the H2B-H4 interface. Next, we demonstrate that these same mutations lower the dimer dissociation temperature and shift the nucleosome dissociation pathway.

## Introduction

The long DNA polymers of eukaryotic cells are perfectly folded in chromosomes, with humans having 23 pairs of chromosomes containing about 3 billion DNA base pairs. The human genome has DNA sequences that code for over twenty thousand proteins. These sequences have been proposed to play a significant role in chromatin dynamics and, in some cases, have been relevant in disease conditions. DNA binding proteins such as polymerases, transcription factors, remodelers, and topoisomerases interact with chromatin to maintain stability and access to cell information. The large-scale dynamics of the chromatin fibers depend on the dynamics of its monomeric unit, the nucleosome core particle (NCP). The NCP (**Figure 1a**) is formed by an octameric protein core containing four types of histones (H3, H4, H2A, and H2B) around which about 146 base pairs of DNA are wrapped (1). At larger structural scales, histones and DNA come together to form chromatin (1–5), which can undergo phase separation to establish distinct functional domains (6,7). The histones are rich in positively charged amino acids. These positively charged amino acids are common sites of post-translational modification (PTMs), such as acetylation, phosphorylation, ubiquitination, and methylation (8). Histone or DNA modifications can change the charge distribution of the NCP, thereby changing its dynamics and altering cellular processes, including transcription, replication, and sometimes DNA repair (9,10).

**Figure 1:**
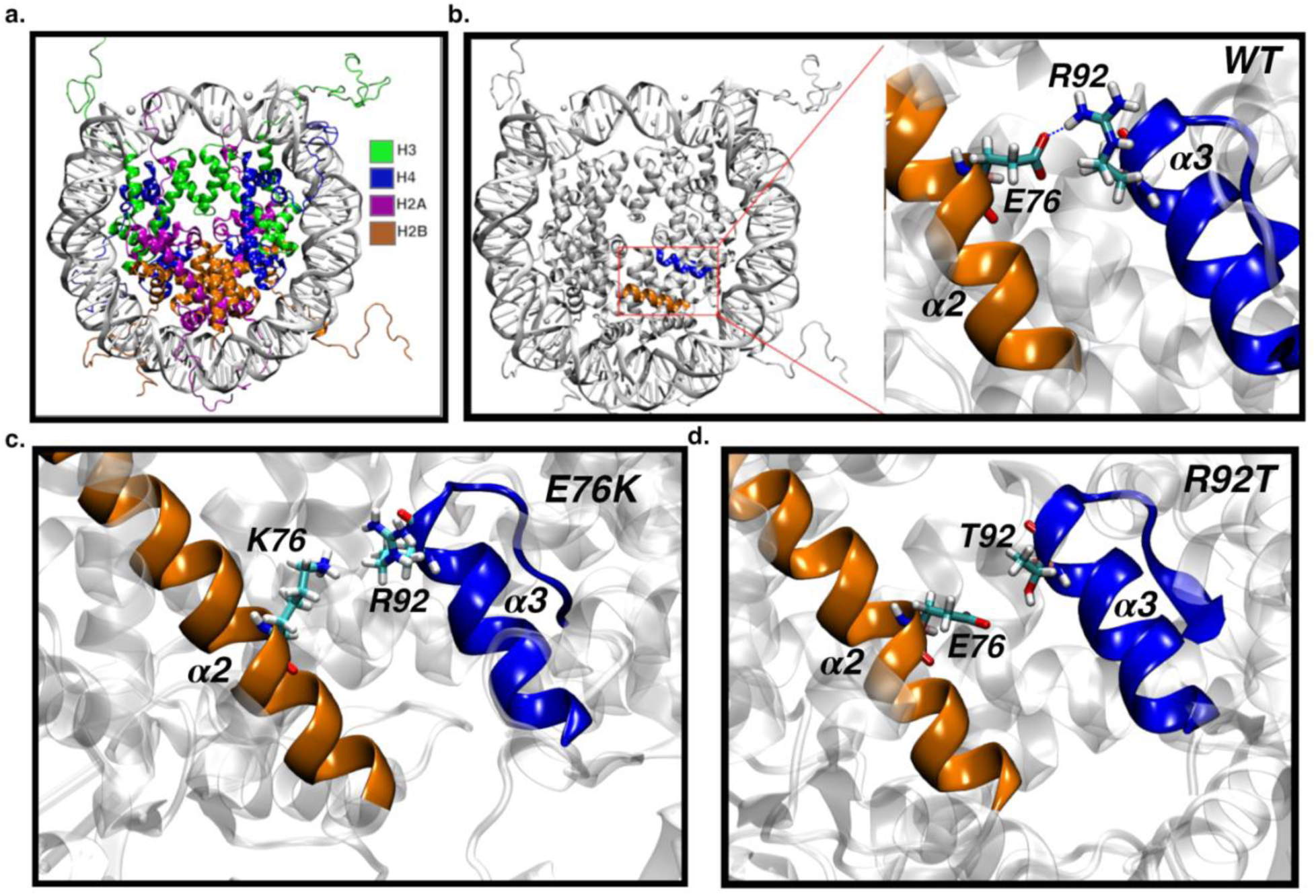
(a) The crystal structure of the nucleosome core particle (NCP) (PDB ID: 1KX5). Histone H2A, H2B, H3 and H4 are colored in magenta, orange, green and blue respectively. (b) The left panel shows the nucleosome highlighting the H4-H2B interface with the orange ⍺2 helix on H2B and the blue ⍺3 helix on H4. The right panel shows the zoomed-in H4-H2B interface with H2BE76 on the ⍺2 helix and H4R92 on the ⍺3 helix. Both amino acids are linked by a salt bridge. (c) The H2BE76K mutation with H2BE76 mutated to H2BK76 (d) The H4R92T mutation with H4R92 mutated to H4T92.

Nucleosome dynamics are associated with an extended range of timescales, such as unwrapping nucleosomes that range from milliseconds to seconds. Small-scale rearrangements, such as breathing or loop formation, happen at much shorter timescales—on the order of microseconds (11–15). Post-translational modifications (PTMs), such as acetylation, methylation, and ubiquitination in both the globular core and disordered tails, can influence the structure and dynamics of the nucleosome (16). For example, mutations have been shown to change the unwrapping rate (17,18). Furthermore, histone variants and histone tail conformations promote DNA bulging and overall dynamic changes of the nucleosome at the microsecond timescale in all atomistic simulations (11,19). This demonstrates that nucleosome-binding proteins elicit stress and deformation in nucleosomes. Histone variants can also have similar effects (20); how these modifications are coupled to nucleosome dynamics when under tension remains an open question. Besides DNA-protein interactions, nucleosome dynamics are affected by the concentration of ions, including Na^+^ and Mg^2+^ (21,22). Chakraborty *et al*., using all atomistic molecular dynamics simulations, showed that the breathing motion seen in DNA during nucleosome unwrapping depends on the concentration of ions (23). Indeed, the motion may also be dependent on the DNA sequence (24). While characterizing the free-energy landscape with coarse-grained simulation, Zhang and colleagues explained that the DNA unwinding and protein disassembly are coupled in the mechanism of nucleosome unfolding (25). Indeed, coarse-grain representations of the nucleosome can aid in characterizing the dynamics of these complexes and their phase behavior (6,26).

Mutations in core histone residues have been seen in various cancer types, including colorectal, lung, breast, and head and neck cancer (27). However, specific mutations in the nucleosome, which drive cellular processes, fundamentally alter these dynamics in cancer cells. Oncogenic mutations in the histone core have been proposed to alter chromatin remodeling processes, such as histone exchange and nucleosome sliding (28). The most frequent mutations are found on both H2B and H3 (27), sites associated with the acidic patch, which are associated with nucleosome-nucleosome interactions, sites on the H3 N-terminal tails, and sites that are *Sin-* mutations, as well as H2AE121 (29) in the H2A C-terminal tail. Most of the above mutations are associated with negatively or positively charged residues; they will disrupt the electrostatic interactions in the NCP primarily through the removal of salt bridges. Here, we focus on characterizing the effect of core histone mutations at the H2B-H4 interface that involve charge reversal or charge removal on the structure of the NCP. Salt bridges between charged amino acid residues contribute to the stability of the histone core. An example of such an interaction is seen at the H4-H2B interface, where the positively charged H4R92 forms a salt bridge with the negatively charged H2BE76 **(Figure 1b)**. Mutations that reverse or nullify these charges have been shown to disrupt the electrostatic interactions that stabilize the histone core. It has been reported that the H2BE76K mutation **(Figure 1c)** destabilizes the H4-H2B helix interface in this manner, causing epigenetic dysregulation (27,28,30). *Arimura* and co-workers resolved the crystal structure of the H2BE76K mutant NCP. They characterized the effect of the mutation using thermal stability assays, fluorescence probes, and gel filtration chromatography to show a decrease in inter-subunit association at the H2A-H2B dimer and the H3-H4 tetramer interface (30). Bagert and colleagues saw that this mutation causes gene dysregulation and loss of transcription silencing (28). However, the mechanism of this destabilization has not been extensively studied at the molecular level using molecular dynamics simulations.

Here, we report long-time molecular dynamics simulations of the effect of the H2BE76K mutation on the nucleosome core particle. H4R92T, a mutation located on a charged arginine across the H2B-H4 interface **(Figure 1d),** is the fifth most probable mutation on histone H4(27), but has not been well characterized. This mutation removes the charge, and we expect an intermediate effect on the stability of the interface. We, therefore, use molecular dynamics (MD) simulations to evaluate the effect of the H2BE76K and the H4R92T on the stability of the H4-H2B interface in the core of the NCP. We hypothesize that the H2BE76K and H4R92T mutations decrease the stability of the H4-H2B interface by increasing the electrostatic repulsion between the H2A/H2B dimer and H3/H4 tetramer. Next, we characterize the effect of the thermodynamic impact of these same mutations on the disassembly of the NCP using differential scanning calorimetry (DSC). We observe differential effects from these two mutations across the H2B-H4 interface.

## Materials and Methods

### NCP Modeling and Simulation

We performed six sets of molecular dynamics simulations of the nucleosome core particle (NCP) for 6 microseconds each, as summarized in **Table 1**. We started from the 1KX5 crystal structure (31). Four of the systems have single-point oncogenic mutations in the histone core at the H2B-H4 interface. These single-point oncogenic mutations of the histone were modeled using *UCSF* Chimera (32). The wild-type (*WT*) system had no mutation, the H2BE76K mutant system had glutamate-76 (Glu76) on histone H2B being substituted with Lysine, while the H4R92T system had arginine-92 (Arg92) replaced with Threonine on histone H4. All mutations were made on one copy of their respective histones. The wild-type and mutant systems above were prepared separately at two different concentrations of Sodium Chloride, 0.15 M physiological conditions and 2.4 *M* high salt concentration. The Mn^2+^ in the crystal structure was substituted with Mg^2+^ All systems were simulated using the AMBER force field (33). Specifically, DNA was simulated using OL15 (34), while the histone was simulated with ff19SB (35). The *OPC* water model was used (36). The Joung and Cheetham parameters for Sodium (Na^+^) and Chlorine (Cl^-^) ions (37) were used. The Lennard Jones parameters between monovalent ions and water were modified following Kulkarni *et al* (38). Mg^2+^ parameters were used from the Li/Merz compromised parameter set (39). Full details regarding the total simulation size are given **Table S1**.

**Table 1:** WT and mutant NCP systems simulated using molecular dynamics.

| System | Wild-Type (WT) | H4R92T | H2BE76K |
| --- | --- | --- | --- |
| Position of Mutation | None | α3 helix on H4 | α2 helix on H2B |
| Residues mutated | None | Arg92 to Thr92 | Glu76 to Lys76 |
| [NaCl] (M) | 0.15 and 2.4 | 0.15 and 2.4 | 0.15 and 2.4 |
| Production run | 6 ms | 6 ms | 6 ms |

Next, we minimized and simulated each system using pmemd in amber18 (40). Systems were minimized using conjugate gradients and steepest descent algorithms for 15 ps each to reduce steric clashes of amino acid sidechains and, hence, reduce stress on the system. Each system was further equilibrated in NVT for 20 ps, as the systems were heated to 310 K (41,42). Subsequently, we used the NPT ensemble for 100 ns with a pressure of 1 bar using the Berendsen barostat (43). The Langevin thermostat was used to maintain the temperature around 310 K with a collision frequency of 1.0 ps (44). The SHAKE algorithm (45) was implemented to freeze hydrogen atoms to allow for a timestep of 2 fs. The Particle Mesh Ewald (PME) (46) algorithm was used to compute the electrostatic interactions in the system, while the Van der Waals cut-off was set to 12 Å. The 1-4 scale was utilized to separate the non-bonded atoms from bonded atoms (47). Next, a 6 µs NPT production run was performed on Anton-2 (48) using a Martyna, Tobias, and Klein (MTK) barostat at a pressure of 1 bar (49,50). Also, the Nose-Hoover thermostat was used to maintain the temperature of the system around 310 K (51). The SHAKE algorithm was implemented to freeze hydrogen atoms to allow for a timestep of 2.5 fs. The short-range bonded and nonbonded forces were computed every time step, while the long-range electrostatic interactions were evaluated once every three time steps. The output energy and trajectory of the system were written every 240 ps.

### Protein expression and purification

#### Protein Expression

Plasmids for the *WT* human core histones were generously supplied by the David laboratory at the Memorial Sloan Kettering Cancer Center in Cornell University. The mutants H2BE76K and H4R92T were generated by site directed mutagenesis using primers (Table S1b). Core histones H2B and H4 were expressed in BL21-DE3 *Escherichia coli (E. coli*) cells while H2A and H3 were expressed in C43 *E. coli* cells. In both cases, cultures were grown until a 600 nm optical density (OD600) of 0.6 - 0.8 was reached, upon which cultures were induced by addition of IPTG to a final concentration of 0.5 mM. Histone H2B and H4 were induced for four hours at 37°C while H2A and H3 were induced overnight at the same temperature. All cells were pelleted, resuspended in Phosphate Buffered Saline (PBS) with 1 mM Phenylmethylsulfonyl fluoride (PMSF) and frozen at −80°C for storage. Pellet resuspensions were thawed and lysed by rod sonication on ice at 60% power with 5 secs on and 10 secs off for 3 minutes. Lysates were cleared by centrifugation at 16,000 ×g and supernatants were discarded. Pellets were resuspended in 6.0 M guanidine in PBS and extracted with mild agitation at 4°C overnight as described by Prescott *et al.* (52). Extracts were cleared by centrifugation at 16,000 ×g and filtered with 0.42 µM syringe filters. An Agilent Zorbax 300SB-C18 semi-preparative High-Performance Liquid Chromatography (HPLC) column attached to an Agilent 1100 series system was equilibrated with 70% HPLC Solvent A (Water with 0.1%TFA) and 30% HPLC Solvent B (Acetonitrile with 0.1% TFA) for 20 minutes. Clean extracts were mixed 1:1 with Solvent A and 0.5 mL of mixture was injected. A gradient from 30-90% Solvent B was run over 30 minutes. Pure histone fractions were identified and evaluated using SDS-PAGE and a Shimadzu MALDI-8020 MALDI-TOF mass spectrometer. Pure fractions of histone were frozen at −80°C and lyophilized for long term storage.

#### DNA purification

A plasmid containing 12 copies of the Widom 601 147 base pair sequence separated by EcoRV restriction digestion sites was provided by the Luger laboratory at the University of Colorado Boulder. The plasmid was transformed into NEB5*⍺ E. coli* cells. Cells were grown and plasmids were extracted using alkaline lysis plasmid isolation and polyethylene glycol 8000 with methods as described by Luger (53). Purified plasmids were digested with EcoRV restriction enzyme and tested on a 1% Agarose gel to verify completion of the digestion. Digested Widom 601 fragments were separated from digested plasmid backbone using polyethylene glycol 8000 with methods as described by Luger (53). The final product was again tested on a 1% Agarose gel to verify a lack of plasmid backbone contamination.

#### Octamer Refolding and NCP Reconstitution

Histone octamer and NCP were made through protocols adapted from the David laboratory (52). Lyophilized core histones were rehydrated in histone unfolding buffer (20 mM Tris-HCl pH 7.6, 6 M guanidine hydrochloride, 1 mM DTT) at 2 mg/mL. Unfolding proceeded for 30 minutes to 3 hours before unfolded histones were centrifuged at 16,000 ×g and concentration of the proteins in the supernatant determined by measuring the absorbance at 280 nm. Histone H3 and H4 were mixed in equimolar quantities while H2A and H2B were added at 5% excess each. The mixture was diluted to 1 mg/mL of total protein concentration before being dialyzed against octamer refolding buffer (10 mM Tris-HCl, pH 7.6, 2 M NaCl, 1 mM EDTA, 1 mM DTT) at 4°C. The dialysis buffer was changed at least three times before the mixture was again centrifuged at 16,000 ×g. The supernatant was concentrated using a 3.5 kDa Amicon centrifugal filter to between 0.5-1.5 mL. The sample was injected 0.5 mL at a time onto a Superdex 200 10/300 GL column attached to an AKTA Go FPLC system (Cytiva). Relevant fractions were collected and tested for the presence of all four core histones using SDS-PAGE and MALDI-TOF mass spectrometry. Fractions containing pure histone octamer were concentrated and stored in 50 % glycerol at −20 °C for future use.

The 147 bp Widom 601 DNA was resuspended in Octamer refolding buffer and combined with *WT* histone octamer in a 1:1.2 molar ratio. The mixture was dialyzed against NCP initial buffer (10 mM phosphate pH 7.6, 1.4 M NaCl, 1 mM EDTA, 5 mM β-mercaptoethanol) before being diluted with dilution buffer (10 mM phosphate pH 7.6, 10 mM NaCl, 1 mM EDTA, 5 mM β-mercaptoethanol) using a peristaltic pump at a flow rate of 0.5 mL/min (52). The product was dialyzed twice against dilution buffer before being centrifuged at 16,000 ×g. Phosphate and β-mercaptoethanol were used for final buffer conditions due to their favorable performance in differential scanning calorimetry (DSC). Mutant NCP (H2BE76K and H4R92T) were prepared by first making H2BE76K dimer or H4R92T tetramer before purifying these species with the same FPLC system mentioned earlier. The DNA, dimer, and tetramer components for each system were then mixed in a 1:1.2:1.2 molar ratio before being subject to the same reconstitution protocol as above. All NCP were analyzed on an electrophoretic mobility shift assay to check for purity and homogeneity using 5% Acrylamide Gels and TBE buffer. Gels were run at 120 V, stained with ethidium bromide and analyzed under UV light.

#### Thermal Stability Assay

Thermal stability of *WT* and mutant NCP samples were tested using an ABI 7500 Real Time PCR system (54,55). NCP was mixed with SYPRO orange (Invitrogen) in dilution buffer to final concentrations of 60 ng/µL DNA and 10× dye (diluted from the 4000× concentration provided by the manufacturer. Thirty microliters of each sample were loaded in triplicate onto 96 well plates. A temperature gradient was run from 26-95°C. Fluorescence was monitored using the VIC channel. Normalized fluorescence intensities were plotted and fit to a sigmoidal curve with the optimize curve fit function in the Python SciPy library using 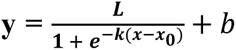. Here y is the normalized fluorescence, x_0_ is the inflection point of the sigmoidal curve corresponding to the melting temperature for the system, *x* is temperature, *L* is the asymptote value of the curve, *k* is growth rate, and *b* is the vertical shift.

#### Differential Scanning Calorimetry (DSC)

For DSC, a TA Nano DSC instrument at the New York University (NYU) Shared Instrument Facility was used (56). NCP samples were concentrated to around 2 mg/mL and 0.6 mL of sample was run against 0.6 mL of dilution buffer. The sample was scanned from 25 to 95 °C at a ramp rate of 1 °C/min. Data was processed and fit using TA Nano Analyze Version 4.0.0.4 data analysis software. A separate run with only buffer against buffer was used for baseline subtraction of the sample run. The run baseline was fit using a polynomial function. Melting temperatures were automatically detected as local maxima. The four peaks were fitted globally using a four event Voigt model. Change in enthalpy for each event was determined as the area under each fitted curve.

#### Binding Free Energy

The Onufriev, Bashford, and Case (OBC) variant of the Generalized Born and Surface Area (MM/GBSA) solvation method (57,58) was used to calculate the binding free energy at the H2B-H4, the tetramer (H3-H4)-dimer (H2A-H2B), and the dimer (H2A-H2B)-DNA interfaces of the histone subunits. The free energy was calculated between the mutated H2B and H4 histone subunits at the H2B-H4 interface, and between H3-H4 tetramer and one or both H2A-H2B dimers. The LEaP command for all systems was set to PBradii mbondi2 (58).The binding free energy ΔG_*bind*_ was calculated (59) ΔG = *ΔG*_*MM*_ + *ΔG*_*sol*_ − *TΔS*. The molecular mechanics contribution ΔG_*MM*_ = ΔE_*int*_ + ΔE_*elec*_ + ΔE_*vdw*_ to the binding free energy included the bond, angle, torsion, electrostatic, and van der Waals terms (60). The bond, angle, and torsion terms were the intramolecular energy contributions, ΔE_*int*_. One trajectory was used to calculate the binding free energy at all interfaces for each system hence the assumption ΔE_*int*_ = 0 as in ΔG_*MM*_ = ΔE_*elec*_ + ΔE_*vdw*_. ΔG_*sol*_ is the solvation free energy contribution having the electrostatic ΔG_*GB*_ and nonpolar solvation energy ΔG_*nps*_ components. The electrostatic component contains the free energy cost between the charge species in the system and the continuum solvent environment. The non-electrostatic component includes the van der Waal’s interaction between solute and solvent molecules and cost of breaking solvent structure around solute, ΔG_*sol*_ = ΔG_*GB*_ + ΔG_*nps*_. The *T*ΔS term is the entropy contribution to the free energy. The entropy contribution was ignored because all the systems were very similar, except for the single point mutation at H2BE76 in the H2BE76K system and H4R92 in the H4R92T system. Therefore, the entropy contributions to the binding free energy were assumed to be approximately equal. The enthalpy of the ligand association with receptor was the difference between the ΔG of complex and the sum of the ΔG of the ligand and receptor as in *ΔG*_*bind*_ = ΔG_*complex*_ – (ΔG_*H*4_ + ΔG_*H*2*B*_). The electrostatic screening effect of monovalent ions was included in the calculation in the same concentration as in each of the systems (0.15 M or 2.4 M). The energy decomposition (energy contribution) of each amino acid to the binding free energy was computed using MM/GBSA in Ambertools (40,61).

#### Hydrogen Bond Analysis

To compute the lifetime of the inter-subunit hydrogen bonds and salt bridges in the histone between the interacting amino acids at the H2B-⍺2 and H4-⍺3 helices, the six systems were run as described above on Anton-2, but the trajectory was saved once every 0.2 ps for 2 ns. The continuous lifetime (62–66) of the histone inter-subunit hydrogen bonds at the H4-H2B interface was calculated using the method of Gowers *et al*. (63) The time autocorrelation function *C_x_(t)* of the hydrogen bonds was calculated, which gives the probability that a hydrogen bond formed between two atoms *i* and *j* at time (t_0_) will be sustained up to time (t) as 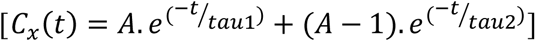. Here, x refers to the definition of lifetime used, *h*_*ij*_(*t*_0_) represents the presence of a hydrogen bond at time t_0_ while *h*_*ij*_(*t*_0_ + *t*) represents the presence of a hydrogen bond after time t. The observed behavior of the hydrogen bond was fitted to a biexponential function using SciPy 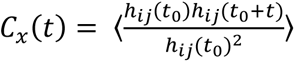 (67) whose integral gives the lifetime of the hydrogen bond(63), 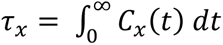.

#### Principal Component Analysis

To characterize the conformations adopted by the helices at the H4-H2B interface, principal component analysis (PCA), a dimensionality reduction technique was applied (68). PCA was performed on the ⍺2 and ⍺3 helices on histones H4 and H2B using coordinates as a collective variable (65,69,70). The interhelical loops were excluded. The combined trajectory of all the systems (*WT* and mutant) at a given NaCl concentration was used. The coordinates of the alpha carbons of the amino acid residues in the above helices at the H4-H2B interface were used to generate a 3N X 3N covariance matrix which was sampled over the combined trajectory (71). Eigenvectors and eigenvalues were obtained from the diagonalized covariance matrix describing the direction of collective motion, and their respective amplitude was used to project the collective variable to the principal components. The first two principal components were used to construct a two-dimensional free energy landscape according to (66,71),*σ*_*ij*_ = 〈(*r*_*i*_ − 〈*r*_*i*_〉) (*r*_*j*_ − 〈*r*_*j*_〉)〉. ΔG(*V*_1_, *V*_2_) = −*k*_*B*_ *T* ln *P* (*V*_1_, *V*_2_) where *V*_1_ and *V*_2_ are the first and second principal components, *P* is the probability density function along *V*_1_ and *V*_2_, *T* is the kelvin temperature and *k*_*B*_ Boltzmann’s constant. The core histone conformations for each system (*WT*, H2BE76K and H4R92T) were extracted from the absolute minima of the PC1-PC2 free-energy plot. Using the Prody normal mode wizard NMWiz (72) in VMD (73), principal component analysis of each system at the minimum was performed (74,75). The eigenvalues and eigenvectors were calculated to describe the magnitude and direction of motion of the ⍺2 and ⍺3 helices in both H2B and H4. The results are visualized through a porcupine plot.

#### Dynamic Cross-Correlation

To evaluate the relative correlated motion of the amino acids in the histone core, we calculated the dynamic cross correlation using Bio3d (76), focusing on histones H4 and H2B. The translational and rotational motions of the protein (77) were removed. The displacement of the histones with time was compared to a reference position, and their displacement was used to calculate the covariance matrix of the atomic weighted mass of the coordinate displacements (78–80). We next determined the fluctuation frequency of the helices and loops at the H4-H2B interface from elastic network modelling for the first three normal modes of each system (76,81).

## Results and Discussion

### Mutation destroys the H2BE76-H4R92 salt bridge at the H4-H2B interface

We perform molecular dynamics simulations on six nucleosome core particle (NCP) systems for 6 microseconds each at two salt concentrations (0.15 M and 2.4 M NaCl). We simulated the wild-type (*WT*) 1KX5 nucleosome and the 1KX5 nucleosome with two oncogenic mutations (H2BE76K and H4R92T). **Figure 2** shows frequent hydrogen bonds at the H2B and H4 interface and their lifetimes. In the wild-type (*WT*) system at 0.15 M, the salt bridge between H2BE76 and H4R92 was the most frequent (**Figure 2a-c**) hydrogen bond between the H4-⍺3 helix and the H2B-⍺2 helix. This salt bridge has an approximate bond distance of 2.0 Å with a lifetime of about 3.3 ps. Also, the H4H75-H2BR92 and phenolic-carbonyl H4Y88-H2BY83 hydrogen bonds are found at the H4-H2B interface, although less frequently. The presence of either the salt bridge between interacting amino acids in the two helices promotes the **π**-**π** interaction between the two aromatic tyrosines (H2BY83 and H4Y88).

**Figure 2:**
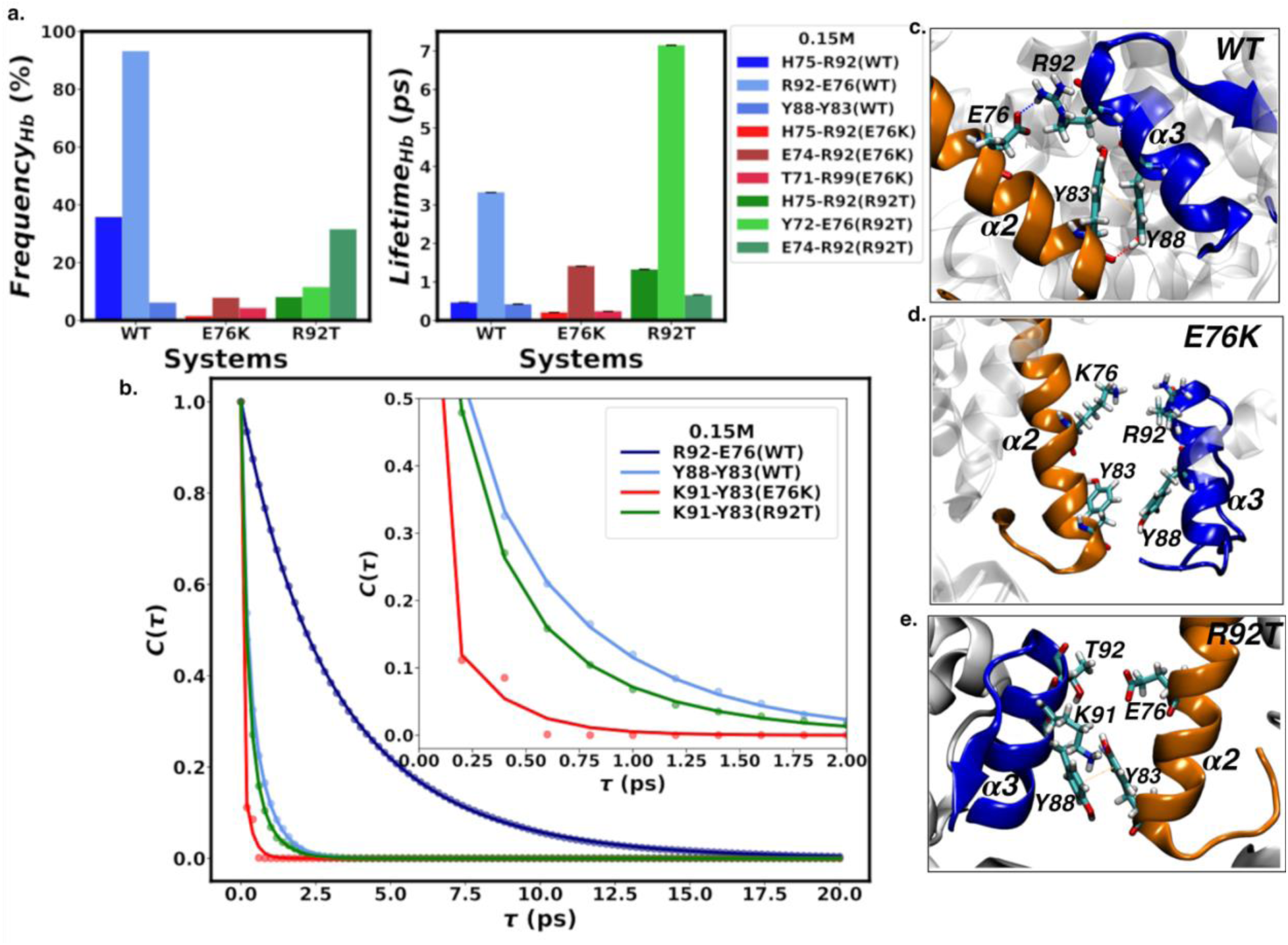
(a) The hydrogen bond frequency and lifetimes of interacting amino acids at the H4-H2B helix interface for WT, H2BE76K and H4R92T systems at 0.15 M NaCl. At the H2B-⍺2 and H4-⍺3 helix interface, the salts bridge between H2BE76 and H4R92 was the most prominent interaction (light-blue) with a 97% frequency and a lifetime of ≈ 3.3 ps. This salt bridge is absent in the mutant systems (H2BE76K and H4R92T). The key shows amino acids labeled in the H4-H2B format. (b) Time autocorrelation function for hydrogen bonds between the H2B-⍺2 and H4-⍺3 helices at the H4-H2B interface. A double exponential curve fit is used to calculate the lifetime using the method of Gowers *et al*., 2015. (c)Amino acid configuration in the *WT* system at 0.15 M showing the salt bridge, hydrogen bonds, and **π**-**π** interaction between interacting amino acids on the orange H2B-⍺2 helix and the blue H4-⍺3 helix (d) The **π**-**π** interaction and salt bridge between the interacting amino acids are broken in the H2BE76K system at 0.15 M (e) The **π**-**π** interaction and some hydrogen bonds are conserved in the H4R92T system, but H4T92 is too short to retain a hydrogen bond with H2BE76.

Conversely, H2BE76K and H4R92T mutations destroy the salt bridge between H2BE76 and H4R92 at the H4-H2B interface at all salt concentrations, as seen by others (82,83). This is consistent with the crystal structure reported by Arimura *et al*. (30), as shown in **Supplementary Figure 1a**. No crystal structure for the H4R92T mutant has been reported. The negative to positive charge reversal in the H2BE76K mutant system pushes the positively charged guanidino group of H4R92 away, leading to a decrease in the frequency of hydrogen bonding between H2BY83 and H4Y88. Consequently, the **π**-**π** H2BY83-H4Y88 interaction decreases as the distance between the two aromatic rings increases (**Figure 2d**). The H4R92T system shows a weaker hydrogen bond between the positively charged ε-amino group of H4K91 and the phenolic group of H2BY83, with a hydrogen bond lifetime of approximately 0.34 ps and a bond distance of 2.0 Å. Like the *WT* system, this hydrogen bond promotes the **π**-**π** association of the two tyrosine rings at this interface (**Figure 2e**). At a high salt concentration of 2.4 M, the hydrogen bond lifetime at the H4-H2B interface was lower compared to physiological salt concentration of 0.15 M. At the same time, the H4Y88-H2BY83 **π**-**π** interaction was conserved for the *WT* and the H4R92T system (**Figure S2**). The H4Y88-H2BY83 **π**-**π** interaction is not conserved for the H2BE76K system.

Overall, the total number of hydrogen bonds between H2B-⍺2 and H4-⍺3 helices at the H4-H2B interface was higher in the *WT* systems when compared to the H2BE76K and H4R92T mutants at all salt concentrations (**Figure S3**). Similarly, the interacting amino acids were closer to each other in the *WT* system when compared with the mutant systems (**Figure S4-S5**). The orientation between the two phenolic rings of H4Y88 and H2BY83 is shown in **Figure S6**.

### Mutations alter the dihedral angle between ⍺2 and ⍺3 helices at the H4-H2B interface

Disruption of the salt bridge between the H2B-⍺2 helix and H4-⍺3 helix in the mutant H2BE76K and H4R92T systems results in a slight shift in the dihedral angle (**θ**) between the helices (**Figure 3a**). The peak in the distribution of the dihedral angle between the plane of the helices in the H2BE76K system is similar to the *WT* system at 0.15 M (**Figure 3b-c).** However, the width of the distribution widens. At 2.4 M (**Figure S7a-b**), there is a shift in the dihedral angle peak for both mutations. The H4R92T system shows a shift in the dihedral angle at 0.15 M and 2.4 M compared to the *WT* (**Figure 3b-c**, **Figure S7a-b**). The slight shift in the hydrogen bond dynamics between H2BE76 and H4R92 upon mutation in the H2BE76K and H4R92T is accompanied by this shift in the dihedral angle between the planes of H2B-⍺2 and H4-⍺3 helices at the H4-H2B interface (**Figure 3d-f**). This shift in the dihedral angle helical plane in the H4R92T system was stabilized by a bifurcated hydrogen bond formed by H4Y72 linking H4T92 and H2BE76 (**Figure 3f**).

**Figure 3:**
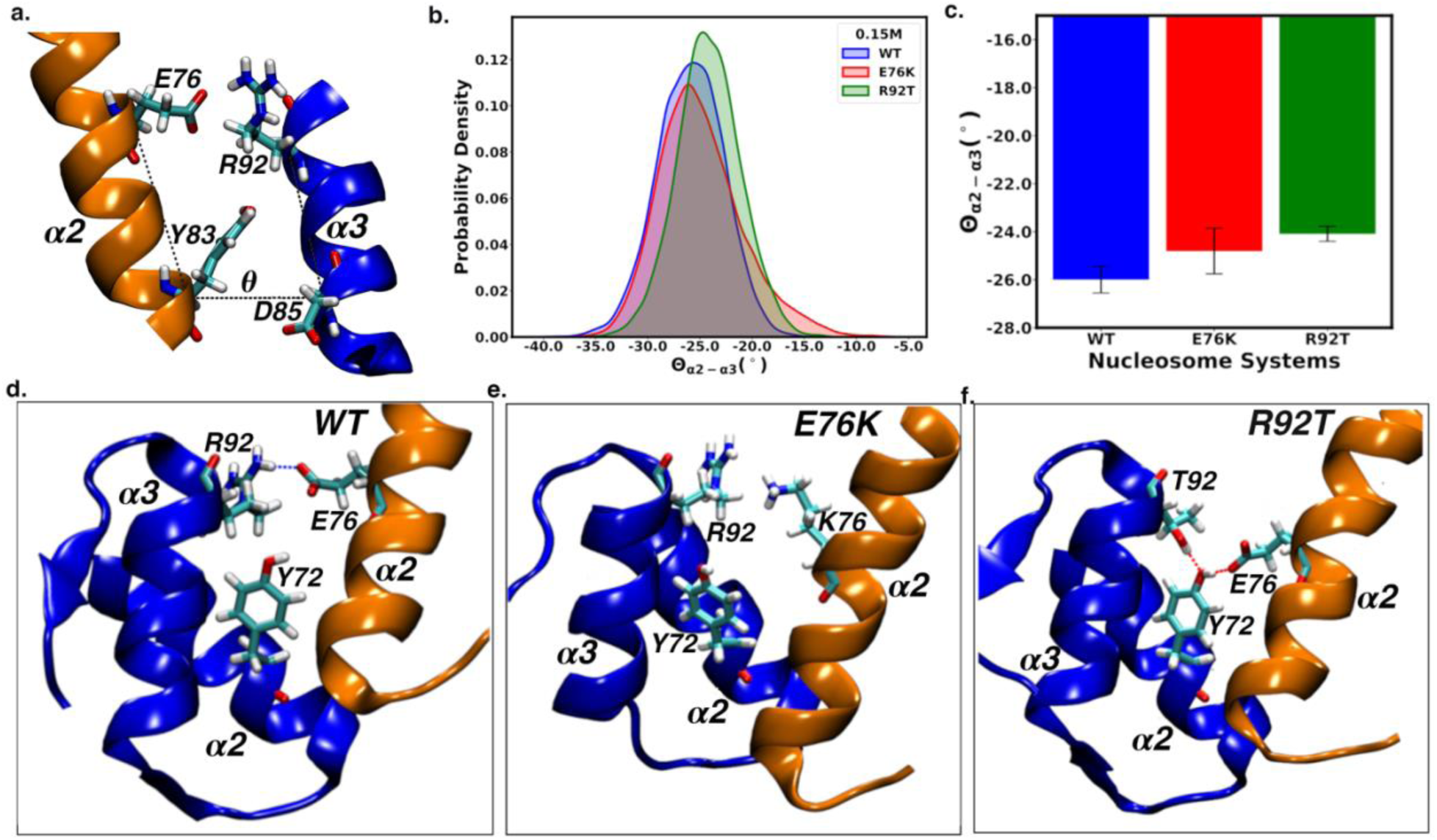
(a) The dihedral angle between the orange H2B-⍺2 helix and the blue H4-⍺3 helix at the H4-H2B interface. The alpha carbon atoms of H2BE76K, H2BY83, H4D85, and H4R92T were used to calculate the dihedral angle between the axes of the helices. (b) Probability density of the dihedral angle and (c) Average and standard error of the mean of the dihedral angle. There is a shift in the helix angle for the H2BE76K and H4R92T systems. The configuration of amino acids that affect the helix dihedral angle for (d) WT, (e) H2BE76K, and (f) H4R92T systems. A bifurcated hydrogen bond linking H2BE76 and H4T92 by H4Y72 is seen in the H4R92T system.

### PCA shows higher interhelical distances in the mutant systems

Principal component analysis (PCA), a dimensionality reduction technique, reveals that the ⍺2 helix of H2B and ⍺3 helix of H4 adopt different helical arrangements in the WT and mutated systems. These helix arrangements differ in the dihedral angle between the plane of the helices and the interhelical distances. **Figure 4a** is the free energy landscape of the first two principal components (PC1, PC2) for the WT and mutated systems combined. The free energy landscape indicates four minima (I, II, III, and IV). The first minimum corresponds to the WT system. Minima II and III correspond to the H2BE76K system. Minima IV corresponds to the H4R92T system. In the *WT* system at 0.15 M *NaCl (Minimum I)*, the helices adopt a conformation with the dihedral angles of −25.0°±2.5° and an average distance of 8.6 ± 0.3 Å between the plane of H2B-⍺2 and H4-⍺3 helices (**Figure 4a-b**). At the same concentration, the interhelical distance in the H2BE76K system (*Minimum III*) was 10.3± 0.4 Å while the H4R92T system (*Minimum IV*) had a 9.4± 0.3 Å separation between the helices (**Figure 4a-b**). The helices drift approximately 1.7 Å apart in the H2BE76K system and approximately 0.3 Å in the H4R92T system compared to the *WT*. The dihedral angles for the H2BE76K system (−24.5°±3.8°) are similar to the WT system, while the H4R92T shows a change from WT in the dihedral helix angle (−21.3°±2.0°) larger than the *WT* and H2BE76K systems (**Figure 4b**). Analysis of the eigenvectors of the first principal component (PC1) and the magnitude (eigenvalues) of PC1 reveal that the H2B-⍺2 and H4-⍺3 helices at the site of mutation in the H2BE76K system have the highest magnitude of motion (**Figure 4c**). Also, the direction of PC1 shows that the helices move away from each other (**Figure 4e**). However, the *WT* and H4R92T systems exhibit a more correlated movement at the H4-H2B helix interface (**Figure 4d-f**).

**Figure 4:**
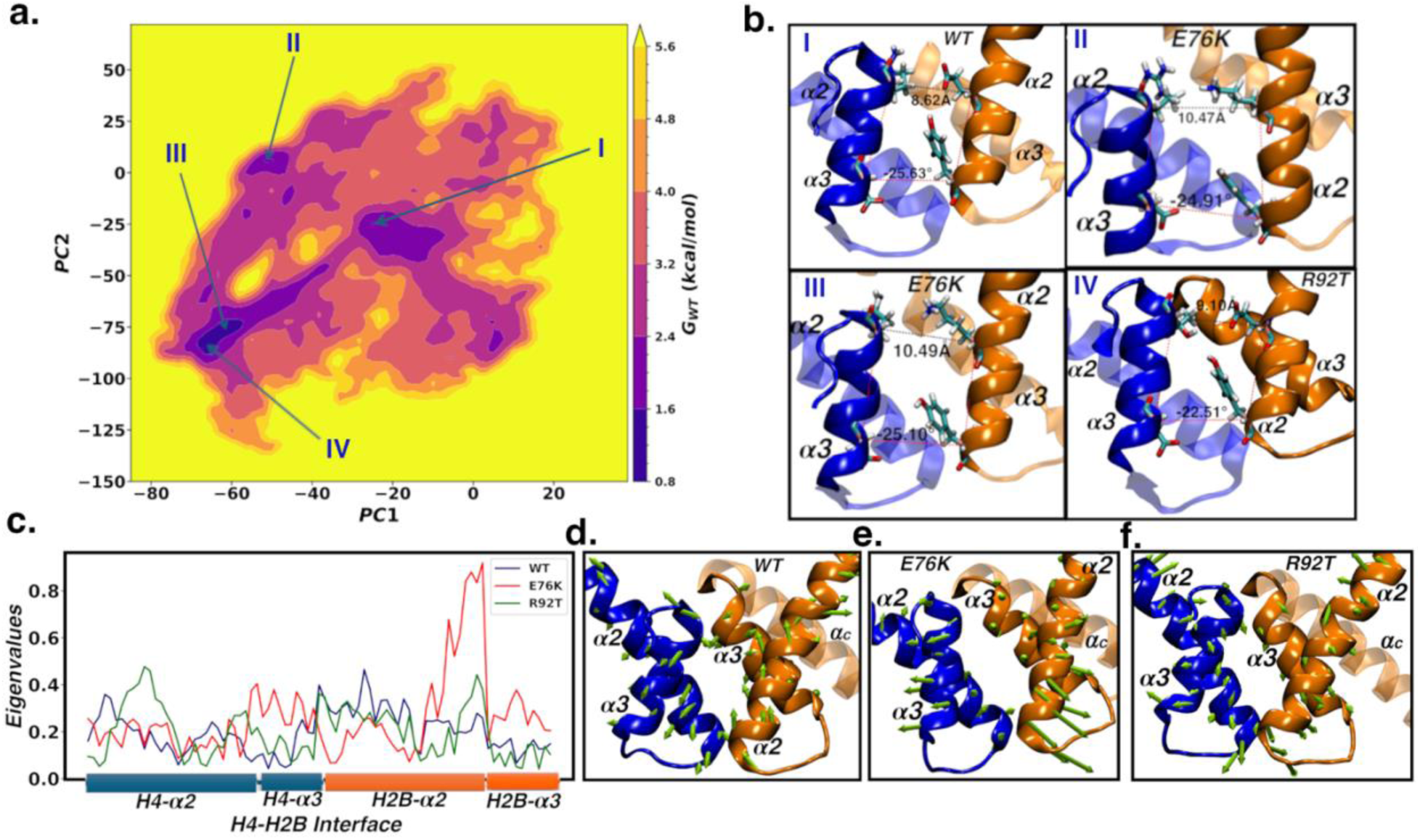
(a) Principal component analysis (PCA) for the combined systems using the coordinates of ⍺2 and ⍺3 helices of both H4 and H2B at the H4-H2B interface at 0.15M of NaCl. The free-energy landscape shows energy minima as I, II, III and IV. (b) The ⍺2 and ⍺3 helix conformations at the H4-H2B interface for WT, H2BE76K and H4R92T systems. The distance and dihedral angles between the plane of the helices are shown. H2BE76K and H4R92T mutations increase the interhelical distance by approximately 1.7 Å and 0.8 Å respectively. (c) Eigenvalues at the H4-H3B interface show H2B-⍺2 and H4-⍺3 in H2BE76K have the highest magnitude of fluctuation. Eigenvector representation of the PC1 for (d) WT, (e) H2BE76K and (f) H4R92T systems. The helices move away from each other in the H2BE76K system, but move towards each other in the WT and H4R92T systems.

The *WT* and mutant systems adopt different helical arrangements at high salt concentrations. The interhelical distance of the *WT* (*Minima I and II*) are statistically similar to the H4R92T system (*Minimum IV*) but are different from the H2BE76K system (*Minimum III*) as shown in **Figure S8a-b**. The interhelical distance in the H2BE76K system increases by approximately 0.7 Å compared to the *WT* (SI text III). The dihedral angle for the H4R92T (−17.9±5.0°) is different from the WT (−22.3±3.0°) and the H2BE76K system (−25.0±1.9°). Like the system at 0.15 M, there is a similar increase in the magnitude (eigenvalues) of motion of the H2B-⍺2 and H4-⍺3 helices in the mutant systems (**Figure S8c-f**)

### Correlated movement of amino acids at the H4-H2B interface decreases with mutation

Dynamic cross correlation between core histone residues reveals that the relative movement of amino acids between H2B-⍺2 and H4-⍺3 helices changes with the mutations at 0.15 M salt (**Figure 5a-c**). There is a higher correlated movement of interacting amino acids at the H4-H2B interface in the WT system than in the mutant systems (H2BE76K and H4R92T), as shown in **Figure 5d-f**. The H2BE76-H4R92 salt bridge and the H2BY83-H4Y88 **π**-**π** interaction promote the correlated movement between the H2B-⍺2 and H4-⍺3 helices as circled in magenta in **Figure 5a**. The H2BE76K system has the least correlated movement of amino acids at this interface as the charge reversal mutation destroys the salt bridge, hydrogen bond, and **π**-**π** interaction of the interacting amino acids (**Figure 5b, 5e**). In the H4R92T mutant system, there is higher correlated motion when compared with the H2BE76K system (**Figure 5c, 5e-f**). The H4R92T mutant preserves the aromatic H2BY83-H4Y88 **π**-**π** interaction and the weaker H4T92-H2BE76 bifurcated hydrogen bond bridged by H4Y72 (**Figure 3f**). These weaker but relevant interactions in the H4R92T system promote the correlated movement of the amino acids at the H2B-⍺2 and H4-⍺3 interface more than the H2BE76K system but less than the *WT* system.

**Figure 5:**
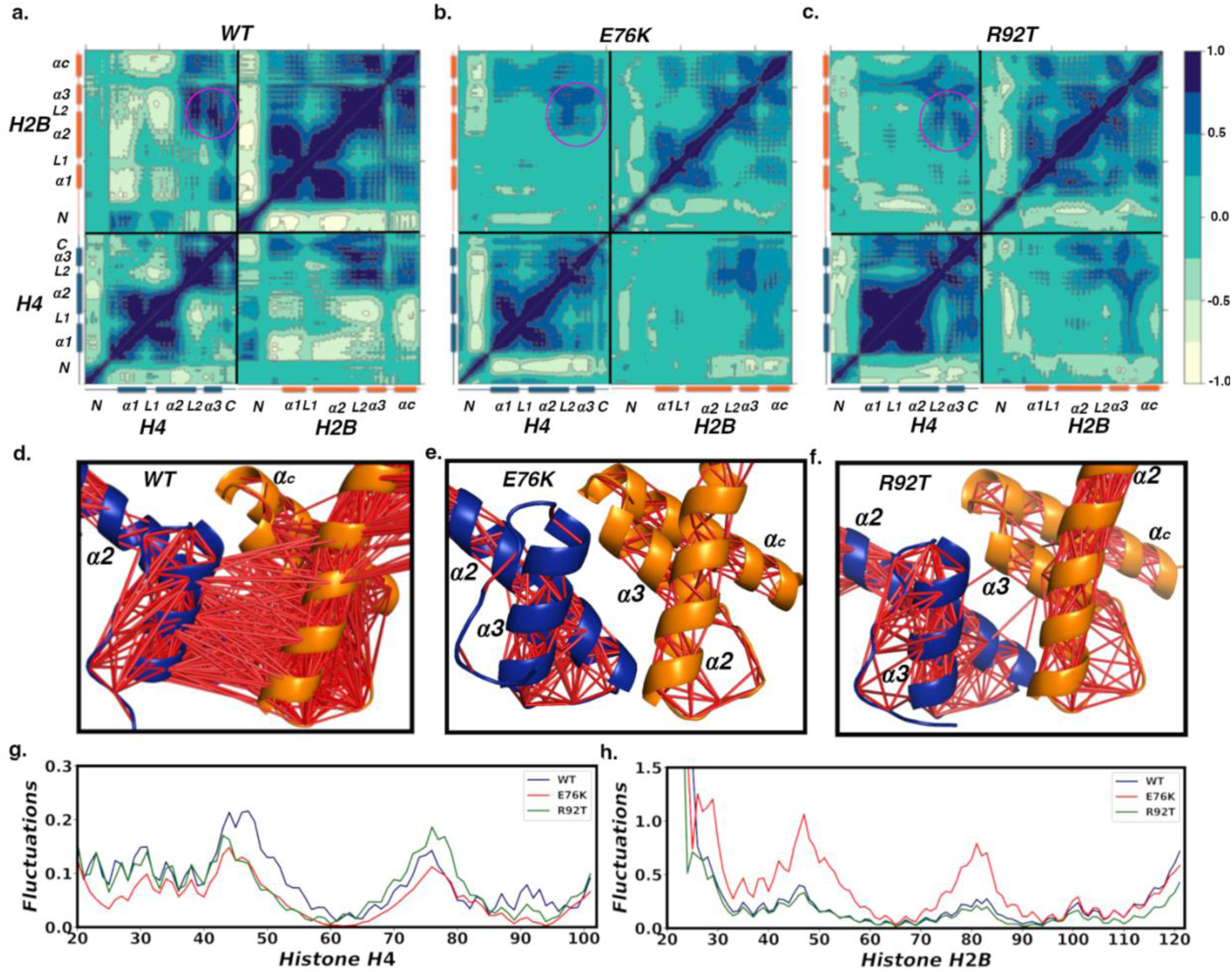
Dynamic Cross-Correlation of the core histone for (a) WT, (b) H2BE76K and (c) H4R92T systems. The mutation changes the overall dynamics of the helices and loops in the histone core. The magenta encircled region is the H4-⍺3 and H2B-⍺2 helix interface. Normal mode analysis ⍺2 and ⍺3 helices at H4-H2B interface showing the magnitude interhelical correlation (d-f). The E76K system has the least correlated motion while the WT system has the most correlated motion. Frequency of motion (fluctuations) at the H4-H2B interface for histone (g) H4 and (h) H2B. Histone H2B in the H2BE76K mutation has the highest fluctuation when compared to other systems. Disruption of hydrogen bonds and salt bridges promote fluctuation.

Analysis of the normal modes shows that the H2BE76K mutation increases the frequency of fluctuation of histone H2B, while that of histone H4 is somewhat similar (**Figure 5g-h**). A decrease in the propensity to form and sustain hydrogen bonds and salt bridges increases the equilibrium fluctuation of helices and loops in the H2BE76K mutant compared to the other systems. Ramaswamy *et al*. showed that the correlated movement of helices and loops in the histone depends on the amino acid sequence of the histone (84).

### Mutation decreases the binding free energy between histone H2B and H4

Molecular mechanics with generalized Born and surface area solvation (MM/GBSA) calculations reveal that the H4-H2B and tetramer (H3-H4)-dimer(H2A-H2B) interfaces in the *WT* system are more stable than the mutant systems at both concentrations (**Figure 6a-c**). Other histone variants have been characterized using similar approaches(85). Locally at the site of mutation (H4-H2B) and globally at the tetramer (H3-H4) and dimer interfaces for one copy (T-D) or both copies (T-DD) of the H2A-H2B dimer(s), the *WT* system is more stable than the H4R92T systems, which in turn is more stable than the H2BE76K systems. The electrostatic contribution to the molecular mechanics component of the binding free energy is positive. In contrast, the electrostatic contribution to the generalized born solvation component of the binding free energy is negative (**Figure S9**). With the H2BE76K mutation, the electrostatic contribution to the molecular mechanics component of the binding free energy increases, while the electrostatic contribution to the generalized born solvation component of the binding free energy becomes more negative. Overall, the mutation results in a less negative binding free energy. Energy decomposition calculations show that the mutation affects the energy contribution of specific amino acids at the ⍺2 and ⍺3 helical interface in both H4 and H2B (**Figure 6d-e**). The global energy contribution of amino acids to the binding at other interfaces besides the H2B-H4 interface are also affected (**Figure S10**). The amino acids H4H75, H4R92, H2BR99, and H2BL100 are the main contributors to the binding free energy at the H4-H2B interface (**Figure S11**). Mutating H2BE76 to H2BK76 at 0.15 M NaCl decreases the residue contribution of H4H75, H4Y88, H4K91, H4R92, H2BK76 (H2BE76 in *WT*), and H2BY83 to the binding free energy. However, the residue contribution of H4T71 and H2BR99 increases (**Figure 6d-e**). There is a significant difference in the free energy of association between a single H2A-H2B dimer and the sugar-phosphate backbone of the double-stranded DNA, as shown in **Table 1**; however, no significant difference in the free energy of association when including both H2A-H2B dimers. This suggests that while the strength of the overall DNA-histone interaction is maintained, the strength of the interaction between the H2A/H2B with the E76K is locally increased. This suggests that DNA accessibility may be more increased more closely to the mutated H2A/H2B.

**Figure 6:**
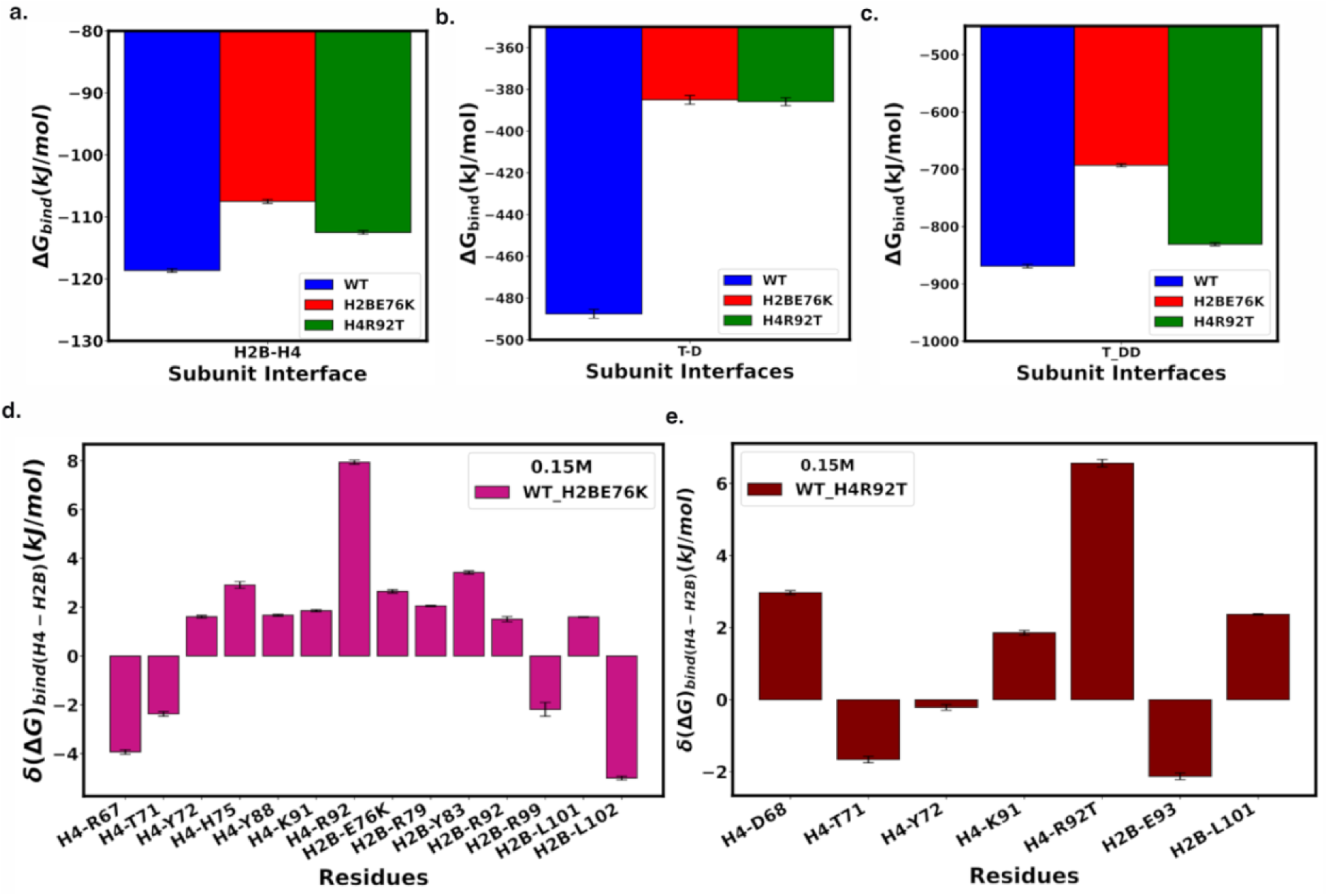
The binding free energy between (a) histone H4 and H2B (b) H3-H4 tetramer and the H2A-H2B dimer with mutant (c) H3-H4 tetramer and both H2A-H2B dimers, including mutant and WT. The *WT* system was more stable than the mutant system at all concentrations. Also, the H4R92T system was more stable than the H2BE76K system. (d) and (e) Variation of the amino acid’s contributions to the binding free energy at the H4-H2B interface upon mutation. Locally at the H2B-H4 interface, the amino acids most affected are at the ⍺2 and ⍺3 helices of H4 and H2B. The amino acid contribution to the binding free energy for H4R92, H4Y88, H2BE76 and H2BY83 decreases with the H2BE76K and H4R92T mutations. The contribution of H4T71 and H2BR99 increases with mutation.

**Table 1:** Free Energy at the Tetramer-Dimer and Dimer-DNA Interface at 0.15 M NaCl (kJ/mol)

| Systems | Nucleosome Interface |  |  |  |
| --- | --- | --- | --- | --- |
|  | T-D | T-DD | D-DNA | DD-DNA |
| **Wild-Type (WT)** | -487.57±2.14 | -868.97±3.17 | -1032.96±6.48 | -2263.94±7.56 |
| **H2BE76K** | -385.03±2.12 | -693.04 ±2.79 | -1205.03±3.55 | -2258.08±7.12 |
| **H4R92T** | -385.92±1.91 | -830.84±2.84 | -1031.33±3.64 | -2129.63±5.00 |

Results reported in *Mean ± SEM*

At a high salt concentration of 2.4 M NaCl, both H2BE76K and H4R92T result in a less negative binding free energy. A similar trend compared to physiological salt was seen in the amino acid interaction at was seen at high salt concentration (**Figure S12**). This suggests that the individual free energy contributions of the interacting amino acids influence the binding free energy at the H2B/H4 interface. This agrees with Arimura and colleagues, who saw that the H2BE76K mutation shifts the melting temperature that correlates with dimer dissociation from the nucleosome core (27,30,82,86). The H4R92T and H2BE76K mutations alter the interaction of the nucleosome with histone-binding proteins, as this mutation occurs spatially near a post-translational modification site (H4K91). The amino acid H4K91 is an acetylation site whose mutation affects the interaction between the NCP and histone acetyltransferases; hence, it mediates DNA protection and chromatin assembly (87,88).

### Core Mutations Differentially Affect Thermal Stability and Enthalpy of NCPs

In complement with our computational work, we desired to probe the stability of the *WT*, H2BE76K, and H4R92T NCP systems using *in vitro* experimental studies. The fluorescence based thermal stability assay (TSA) developed by *Arimura et al*. can determine the relative thermal stability of NCP mutants (28,30). In this assay, a fluorescent dye (SYPRO Orange™) is mixed with NCP in low salt buffer. When the dye binds to hydrophobic regions of proteins, it fluoresces with excitation/emission wavelengths of 470/570 nm (89). While the histones are contained in the intact NCP, these binding sites are unavailable. While the complex dissociates these regions are exposed and the dye can bind to the free histones, increasing the fluorescence signal. Using a quantitative PCR (qPCR) system, the temperature is increased uniformly, and fluorescence is monitored as the NCP dissociates, with higher fluorescence representing an increase in free protein in solution due to dissociation. This process occurs in two stages. First, the dimer is released at a lower temperature, followed by the tetramer release at a higher temperature due to its higher affinity for DNA (**Figure 7a**). Histone mutations, which affect the stability of the NCP, have a correlated effect on the melting temperatures of dimer dissociation. We performed TSA on the *WT*, H2BE76K, and H4R92T NCP systems to assess the relative stability of these complexes and compare these results with our computational findings. Our NCP samples were prepared using recombinant human core histones (or human mutants) and Widom-601 147 bp DNA. In each of these cases, we see the expected biphasic dissociation process of the dimer and tetramer from the DNA. *WT* NCP displayed the highest H2A/H2B dimer melting temperature of 71.5 °C. E76K NCP showed the lowest H2A/H2B dimer melting temperature at 66.3 °C. H4R92T NCP showed an intermediate dimer melting temperature of 70.3 °C (**Figure 7b-e**). The strong destabilization of E76K and intermediate destabilization of R92T NCP, compared to WT, are in line with our MM/GBSA calculations of the binding free energy between dimer/tetramer, as shown in **Figure 6b**. These results also provide us with reliable values for the dimer and tetramer melting temperatures, which prove essential for our next set of experiments.

**Fig 7:**
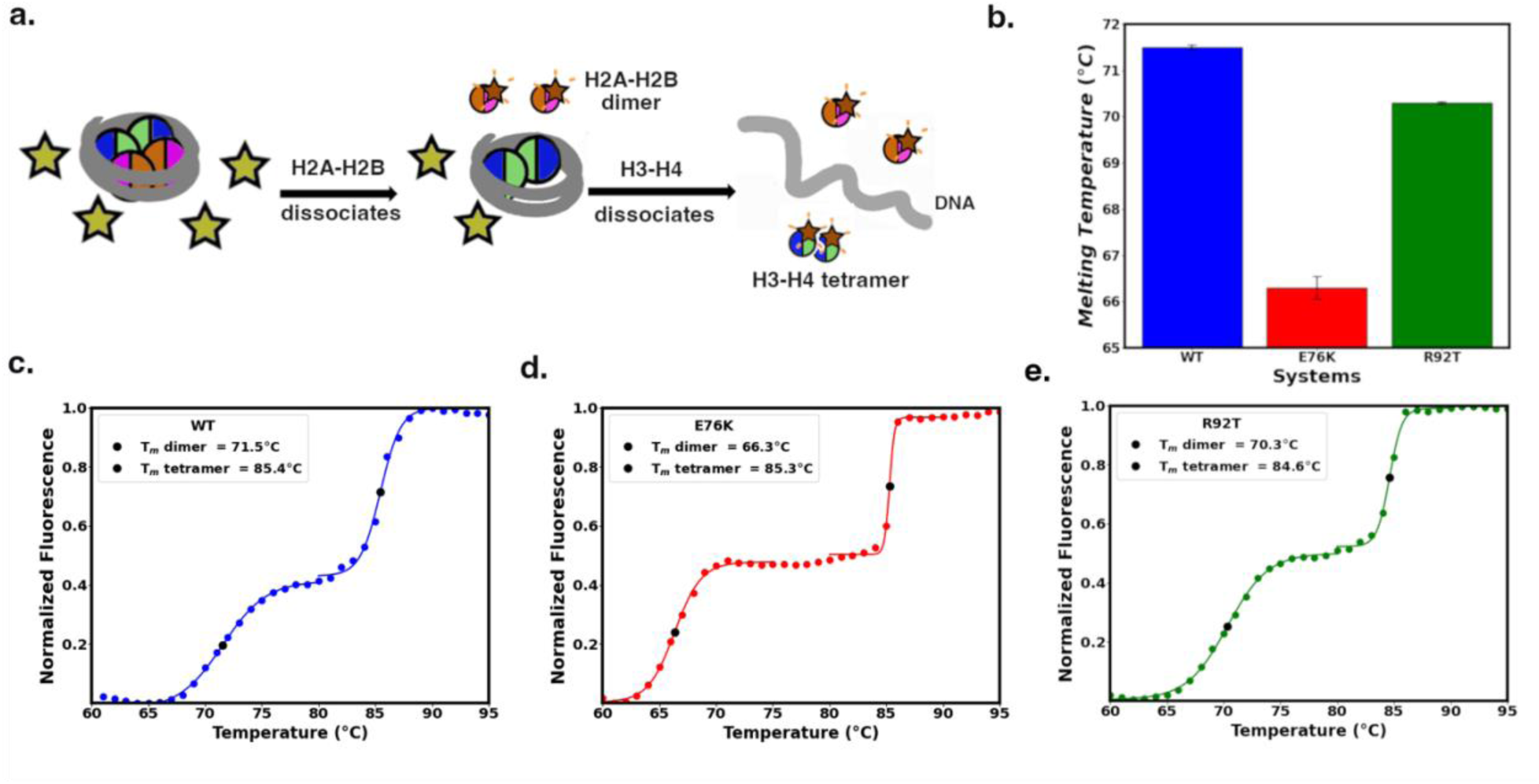
(a) Schematic illustrating the biphasic melting of the nucleosome core particle (NCP) resulting in the binding of the SYPRO Orange™ dye and subsequent increase in fluorescence. (b) Bar chart comparison for the dimer melting temperatures of the three systems as determined by TSA (c) *WT* NCP (d) H2BE76K NCP and (e) H4R92T NCP TSA data with normalized fluorescence plotted against temperature. Each plot demonstrates two sigmoidal curves, where the first represents H2A-H2B dimer dissociation and the second represents H3-H4 tetramer dissociation.

**Figure 8:**
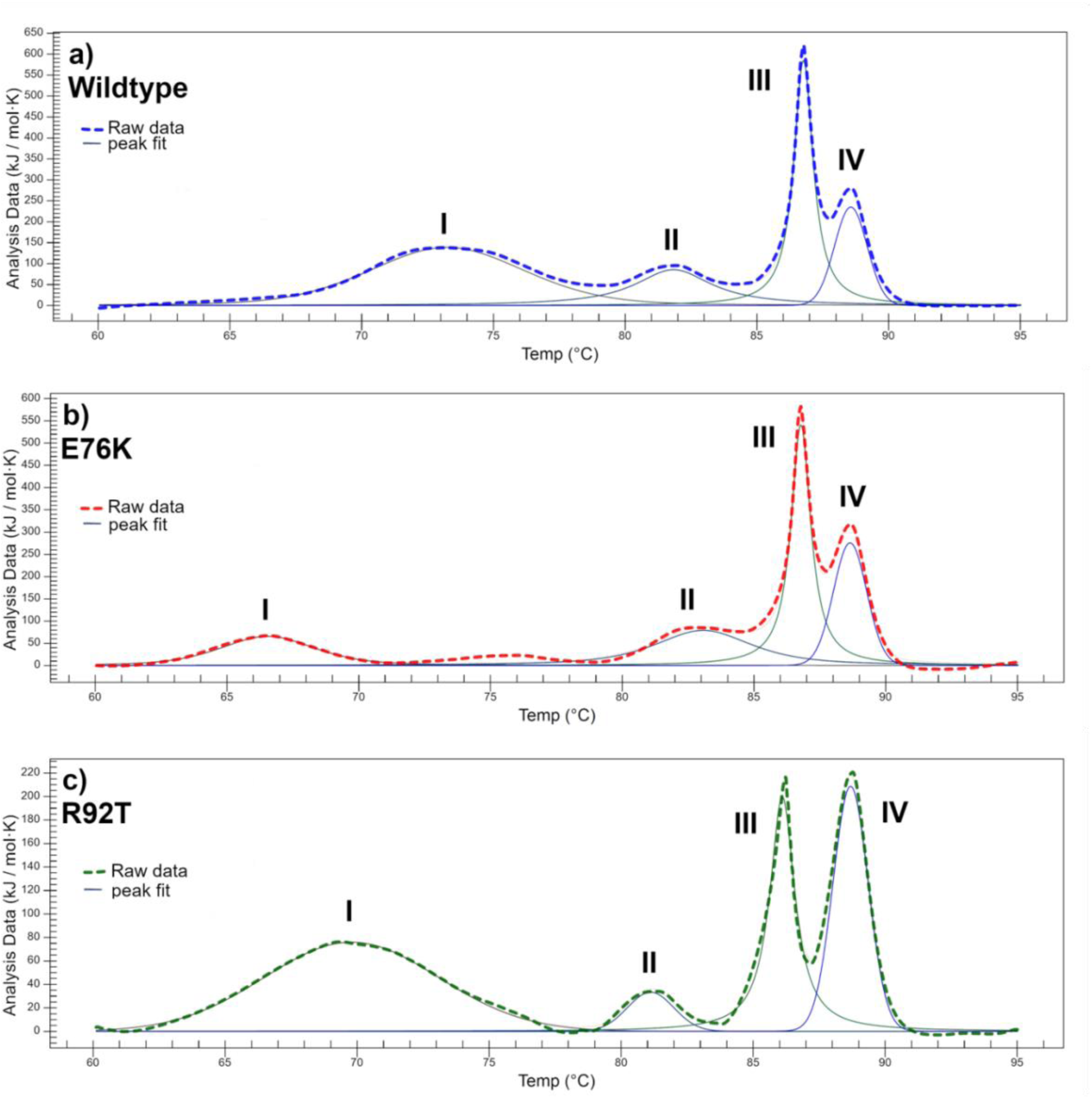
Differential scanning calorimetry of (a) WT, (b) H2BE76K, and (c) H4R92T nucleosome core particle. Peaks I and III correspond to H2A-H2B dimer and H3-H4 tetramer dissociation, respectively. Melting temperature and enthalpy (ΔH) for the H2A-H2B dimers for *WT* are higher than for all mutant systems.

To further explore the effects of these mutations, we perform differential scanning calorimetry (DSC) on our NCP systems. In DSC, the difference in heat flow required for even heating of the sample and a buffer only reference cell is measured. If an event with an associated enthalpy change occurs (such as the breaking of intermolecular bonds in a complex), the heat flow to the sample cell will differ from that of the reference cell. This difference, plotted as a function of temperature, appears as a peak corresponding to the thermal transition. The local maximum of this peak represents the melting temperature (T_m_) for this event. Converting the heat flow into molar heat capacity and integrating over the peak provides the enthalpy change (ΔH) associated with the event.

Previously, Bina *et al.* performed DSC on isolated NCPs under varying salt conditions and compared with free DNA, suggesting that the complexation of histone with DNA is driven primarily by the entropic stabilization of DNA (90). This study found three peaks in the range from 45 – 95 °C. Several studies in the 1980s and 1990s used DSC to study the thermal denaturation of nucleosomes and chromatin derived from animal tissue (91–93). The breakdown of the complex was found to be a multistep process made up of distinct, resolvable events between 45-95 °C, which are highly sensitive to the ionic strength of the solution. More recently, Kolomijtseva *et al.* used DSC to characterize chromatin melting in animal nuclei and found that the concentration of Mg^2+^ ions can shift the melting profile and affects the separation of the three resolvable peaks (91). This study used rat hepatocyte-derived chromatin and found that the two main observable peaks had the highest resolution in 1 mM EDTA. However, the identity of these peaks was not completely clear. The tissue-derived nature of the samples in the studies listed above means that the chromatin contains heterogeneous DNA sequences. It is possible that differences in sequence affinity for the histones could reduce resolution in resultant thermograms. We perform the DSC studies of recombinant NCPs and identify four resolvable peaks for our WT and mutant systems. Additionally, by comparing with our TSA data, we can assign the identity of these peaks.

By comparing the melting temperature values of our DSC and TSA results, we identify DSC peaks I and III as the dimer and tetramer dissociation, respectively. The characteristics of dimer dissociation are of great interest to this study. The WT NCP system has the highest ΔH for the dimer dissociation at 1105 kJ/mol. The E76K NCP system has a ΔH value of 31% of the WT NCP value at 344.0 kJ/mol. While the R92T NCP system’s ΔH value is 61% of the WT NCP value at 672 kJ/mol (**Table 2**). The reduction in ΔH between the WT and E76K NCP systems reflects the high levels of destabilization caused by the repulsion of K76 and R92, as seen in our simulations. The intermediate reduction in ΔH between WT and R92T NCP can likely be attributed to the compensatory intermolecular forces we have identified in R92T, but not E76K NCP. These results vindicate our hypotheses about the stability of these systems and indicate that dimer-tetramer interactions drive the destabilization in these mutations.

**Table 2:** DSC Melting temperatures and enthalpy changes for all systems.

| Systems | DSC Dissociation Peaks |  |  |  |  |  |  |  |
| --- | --- | --- | --- | --- | --- | --- | --- | --- |
|  | I |  | II |  | III |  | IV |  |
| | Temp<br>(°C) | $\Delta H$<br>(kJ/mol) | Temp<br>(°C) | $\Delta H$<br>(kJ/mol) | Temp<br>(°C) | $\Delta H$<br>(kJ/mol) | Temp<br>(°C) | $\Delta H$<br>(kJ/mol) |
| **WT** | 73.2 | 1105 | 81.8 | 453 | 86.8 | 790 | 88.6 | 366 |
| **H2BE76K** | 66.5 | 344 | 83.1 | 473 | 86.8 | 744 | 88.7 | 440 |
| **H4R92T** | 69.8 | 672 | 81.1 | 73.7 | 86.1 | 346 | 88.7 | 350 |

A comparison of the values for tetramer dissociation shows that WT and E76K NCP have similar values at 790 and 744 kJ/mol, respectively. This is expected since both systems contain WT tetramer. Interestingly, the R92T NCP had a ΔH of 346 kJ/mol, a marked 56% reduction in comparison with WT NCP. We can infer that conformational changes seen in the R92T tetramer affect the complex’s affinity for DNA. We suggest that peak II is a rearrangement of DNA around the tetramer after the dimer has dissociated. This is supported by the similar values of ΔH seen for peak II in the WT and E76K systems, since both contain WT tetramer. However, we see significantly lower values of ΔH for this peak in the R92T NCP system. It is likely that conformational changes in the tetramer driven by this mutation prevent the reorganization seen in the other systems. We suggest that peak IV is the denaturation of the double-stranded DNA. This is evidenced by the similar melting temperatures and ΔH values for all three systems.

## Conclusion

Here, we report molecular dynamics simulation to explore H2BE76K and H4R92T oncogenic mutations in the histone core of the NCP at different salt concentrations and predict how these mutations affect the hydrogen bond dynamics of core amino acid residues. We performed multiple microseconds-long all-atomistic simulations of the *WT* and mutant (H4R92T and H2BE76K) NCP systems at two different concentrations of NaCl (0.15 M and 2.4 M). We observe that H2BE76K and H4R92T mutations destabilize the H4-H2B interface in the core of the NCP to various degrees. At all salt concentrations, these mutations weaken the electrostatic interaction between amino acids in the H2B-⍺2 and H4-⍺3 helices. Shifts in hydrogen bond dynamics characterize this destabilization. As expected, with the H2BE76K charge-reversal mutation, the electrostatic contribution to the binding free energy between the H2A/H2B dimer and H3/H4 tetramer becomes more repulsive at all salt concentrations. The H4R92T system has an intermediate dimer-tetramer binding free energy between the H2BE76K system and WT. Thus, we hypothesize that the H2BE76K and H4R92T mutations may affect nucleosome assembly and disassembly thermodynamics. Upon mutation, the weakened electrostatic interaction at the H4-H2B interface will favor nucleosome disassembly with the H2A-H2B dimer dissociation at lower temperatures. This is consistent with the mechanism of increased dimer exchange (28).

Our TSA and DSC results agree with and corroborate our computational findings. We show that the H4R92T NCP has an intermediate H2A/H2B dimer dissociation temperature between *WT* and H2BE76K NCP. Destabilization of the histone core may also alter how nucleosome binding proteins interact with and regulate the dynamics of the NCP *in vitro* and *in vivo* (94). The stability of the helices at higher temperatures with mutations may decrease. We can hypothesize that this is why the melting temperature shifts. The increase in histone DNA binding free energy that is centered on the H2A/H2B with the mutation suggests that the DNA may be less accessible. This may also modulate the binding with transcription factors or chromatin remodelers. While numerous mutations have been mapped, very few oncogenic mutations and their impact on the structure of the NCP and its assemblies have been probed using structural methods (95). Very few oncogenic mutations have been probed computationally (96). The impact of this work is to reveal how distinct sites in neighboring regions of the histone core can lead to similar structural modifications at key protein-protein interfaces, yet functionally different modifications in the dissociation pathway. We hypothesize that these same mutations may modulate the accessibility of the DNA through the expansion of the histone core. Gene expression characterized by chromatin accessibility can also increase in the mutant systems because the destabilized histone core may promote stress in the double helix DNA, which affects DNA repositioning and, in turn, can alter gene expression (82). Nucleosome assembly and disassembly are essential in maintaining cellular homeostasis and allowing the continuous responsiveness of the nucleosome to external and cellular stimuli (97). The assembly and disassembly of the NCP is characterized by histone exchange aided by chaperones, allowing RNA polymerase II (Pol II) to access the open reading frames (ORFs) of genes (98–101). Pol II can also access the transcribable DNA portion of nucleosome when the double-stranded DNA slides over the octameric histone, especially in moderately transcribed genes (97). In all of these processes, reassembly of the nucleosome is needed to prevent cryptic transcription initiation by Pol II, which may synthesize deleterious truncated proteins (101). We suggest that reassembly in the oncogenic mutants is also inhibited because their stability is reduced and that this could be another mechanism for the mutations’ oncogenic effects.

## Data and Code Availability Statement

Analysis codes are available on https://github.com/CUNY-CSI-Loverde-Laboratory/Molecular_Dynamics_of_Structural_Effect_of_Oncogenic_Mutations. Trajectories are available on Zenodo.

## Author Contributions

A.O. performed simulations and analysis. R.P. helped with analysis. C.F. reconstituted the nucleosomes, did TSA and nano-DSC. S.F. and S.M.L. supervised the project.

## Acknowledgements

NIH supported this work through Grant 1R15GM146228-01. Anton 2 computer time was provided by the Pittsburgh Supercomputing Center (PSC) through Grant R01GM116961 from the National Institutes of Health. The Anton 2 machine at PSC was generously made available by D.E. Shaw Research. A. O. would also like the NSF MolSSI and CUNY Dissertation fellowship for funding. We are grateful to Prof. Yael David for discussion and to Qingzeng Gao and Dr. Chin H. Lin for their help. We thank Prof. Chang-Hui Shen for use of the qPCR instrument. A. O. is grateful for the help from Dr. Phu Tang.

## References

1. Kornberg, R. D. 1974. Chromatin structure: a repeating unit of histones and DNA. Science. 184(4139):868–871, doi: 10.1126/science.184.4139.868.

2. McGinty, R. K., and S. Tan. 2015. Nucleosome Structure and Function. Chemical Reviews. 115(6):2255–2273, doi: 10.1021/cr500373h, 10.1021/cr500373h.

3. Segal, E., Y. Fondufe-Mittendorf, L. Chen, A. Thåström, Y. Field, I. K. Moore, J. P. Wang, and J. Widom. 2006. A genomic code for nucleosome positioning. Nature. 442(7104):772–778, doi: 10.1038/nature04979.

4. Luger, K., A. W. Mäder, R. K. Richmond, D. F. Sargent, and T. J. Richmond. 1997. Crystal structure of the nucleosome core particle at 2.8 A resolution. Nature. 389(6648):251–260, doi: 10.1038/38444.

5. Huertas, J., and V. Cojocaru. 2021. Breaths, twists, and turns of atomistic nucleosomes. Journal of molecular biology. 433(6):166744.

6. Farr, S. E., E. J. Woods, J. A. Joseph, A. Garaizar, and R. Collepardo-Guevara. 2021. Nucleosome plasticity is a critical element of chromatin liquid–liquid phase separation and multivalent nucleosome interactions. Nature communications. 12(1):2883.

7. Elathram, N., B. E. Ackermann, E. T. Clark, S. R. Dunn, and G. T. Debelouchina. 2023. Phosphorylated HP1α-Nucleosome Interactions in Phase Separated Environments. Journal of the American Chemical Society. 145(44):23994–24004, doi: 10.1021/jacs.3c06481, <GO to ISI>://WOS:001092726200001.

8. Bowman, G. D., and M. G. Poirier. 2015. Post-Translational Modifications of Histones That Influence Nucleosome Dynamics. Chemical Reviews. 115(6):2274–2295, doi: 10.1021/cr500350x, 10.1021/cr500350x.

9. Fenley, A. T., D. A. Adams, and A. V. Onufriev. 2010. Charge State of the Globular Histone Core Controls Stability of the Nucleosome. Biophysical Journal. 99(5):1577–1585, doi: 10.1016/j.bpj.2010.06.046, 10.1016/j.bpj.2010.06.046.

10. Ngo, T. T. M., J. Yoo, Q. Dai, Q. Zhang, C. He, A. Aksimentiev, and T. Ha. 2016. Effects of cytosine modifications on DNA flexibility and nucleosome mechanical stability. Nature Communications. 7(1):10813, doi: 10.1038/ncomms10813, 10.1038/ncomms10813.

11. Shaytan, A. K., G. A. Armeev, A. Goncearenco, V. B. Zhurkin, D. Landsman, and A. R. Panchenko. 2016. Coupling between histone conformations and DNA geometry in nucleosomes on a microsecond timescale: atomistic insights into nucleosome functions. Journal of molecular biology. 428(1):221–237.

12. Li, Z., and H. Kono. 2016. Distinct roles of histone H3 and H2A tails in nucleosome stability. Scientific reports. 6(1):31437.

13. Gansen, A., F. Hauger, K. Toth, and J. Langowski. 2007. Single-pair fluorescence resonance energy transfer of nucleosomes in free diffusion: optimizing stability and resolution of subpopulations. Analytical biochemistry. 368(2):193–204.

14. Chen, Y., J. M. Tokuda, T. Topping, S. P. Meisburger, S. A. Pabit, L. M. Gloss, and L. Pollack. 2017. Asymmetric unwrapping of nucleosomal DNA propagates asymmetric opening and dissociation of the histone core. Proceedings of the National Academy of Sciences. 114(2):334–339.

15. Ngo, T. T., Q. Zhang, R. Zhou, J. G. Yodh, and T. Ha. 2015. Asymmetric unwrapping of nucleosomes under tension directed by DNA local flexibility. Cell. 160(6):1135–1144.

16. Bowman, G. D., and M. G. Poirier. 2014. Post-translational modifications of histones that influence nucleosome dynamics. Chemical reviews. 115(6):2274–2295.

17. Simon, M., J. A. North, J. C. Shimko, R. A. Forties, M. B. Ferdinand, M. Manohar, M. Zhang, R. Fishel, J. J. Ottesen, and M. G. Poirier. 2011. Histone fold modifications control nucleosome unwrapping and disassembly. Proceedings of the National Academy of Sciences. 108(31):12711–12716.

18. Kim, J., J. Lee, and T.-H. Lee. 2015. Lysine Acetylation Facilitates Spontaneous DNA Dynamics in the Nucleosome. The Journal of Physical Chemistry B. 119(48):15001–15005, doi: 10.1021/acs.jpcb.5b09734, 10.1021/acs.jpcb.5b09734.

19. Shaytan, A. K., D. Landsman, and A. R. Panchenko. 2015. Nucleosome adaptability conferred by sequence and structural variations in histone H2A-H2B dimers. Current Opinion in Structural Biology. 32:48–57, doi: 10.1016/j.sbi.2015.02.004, <GO to ISI>://WOS:000359032300009.

20. Wen, Z., L. Zhang, H. Ruan, and G. Li. 2020. Histone variant H2A. Z regulates nucleosome unwrapping and CTCF binding in mouse ES cells. Nucleic acids research. 48(11):5939–5952.

21. Andreeva, T. V., N. V. Maluchenko, A. L. Sivkina, O. V. Chertkov, M. E. Valieva, E. Y. Kotova, M. P. Kirpichnikov, V. M. Studitsky, and A. V. Feofanov. 2022. Na+ and K+ Ions Differently Affect Nucleosome Structure, Stability, and Interactions with Proteins. Microscopy and Microanalysis. 28(1):243–253, doi: 10.1017/s1431927621013751, <GO to ISI>://WOS:000757458900022.

22. Chakraborty, K., M. Kang, and S. M. Loverde. 2018. Molecular Mechanism for the Role of the H2A and H2B Histone Tails in Nucleosome Repositioning. Journal of Physical Chemistry B. 122(50):11827–11840, doi: 10.1021/acs.jpcb.8b07881, <GO to ISI>://WOS:000454566600002.

23. Chakraborty, K., and S. M. Loverde. 2017. Asymmetric breathing motions of nucleosomal DNA and the role of histone tails. The Journal of Chemical Physics. 147(6).

24. Khatua, P., P. K. Tang, A. Ghosh Moulick, R. Patel, A. Manandhar, and S. M. Loverde. 2024. Sequence Dependence in Nucleosome Dynamics. The Journal of Physical Chemistry B. 128(13):3090–3101, doi: 10.1021/acs.jpcb.3c07363, 10.1021/acs.jpcb.3c07363.

25. Zhang, B., W. Zheng, G. A. Papoian, and P. G. Wolynes. 2016. Exploring the Free Energy Landscape of Nucleosomes. Journal of the American Chemical Society. 138(26):8126–8133, doi: 10.1021/jacs.6b02893, 10.1021/jacs.6b02893.

26. Ozer, G., A. Luque, and T. Schlick. 2015. The chromatin fiber: multiscale problems and approaches. Current Opinion in Structural Biology. 31:124–139, doi: 10.1016/j.sbi.2015.04.002, <GO to ISI>://WOS:000357706000018.

27. Nacev, B. A., L. Feng, J. D. Bagert, A. E. Lemiesz, J. Gao, A. A. Soshnev, R. Kundra, N. Schultz, T. W. Muir, and C. D. Allis. 2019. The expanding landscape of ‘oncohistone’ mutations in human cancers. Nature. 567(7749):473–478, doi: 10.1038/s41586-019-1038-1, 10.1038/s41586-019-1038-1.

28. Bagert, J. D., M. M. Mitchener, A. L. Patriotis, B. E. Dul, F. Wojcik, B. A. Nacev, L. J. Feng, C. D. Allis, and T. W. Muir. 2021. Oncohistone mutations enhance chromatin remodeling and alter cell fates. Nature Chemical Biology. 17(4):403-+, doi: 10.1038/s41589-021-00738-1, <GO to ISI>://WOS:000624704100001.

29. Chew, G.-L., M. Bleakley, R. K. Bradley, H. S. Malik, S. Henikoff, A. Molaro, and J. Sarthy. 2021. Short H2A histone variants are expressed in cancer. Nature communications. 12(1):490.

30. Arimura, Y., M. Ikura, R. Fujita, M. Noda, W. Kobayashi, N. Horikoshi, J. Y. Sun, L. Shi, M. Kusakabe, M. Harata, Y. Ohkawa, S. Tashiro, H. Kimura, T. Ikura, and H. Kurumizaka. 2018. Cancer-associated mutations of histones H2B, H3.1 and H2AZ1 affect the structure and stability of the nucleosome. Nucleic Acids Research. 46(19):10007–10018, doi: 10.1093/nar/gky661, <GO to ISI>://WOS:000450955800015.

31. Davey, C. A., D. F. Sargent, K. Luger, A. W. Maeder, and T. J. Richmond. 2002. Solvent Mediated Interactions in the Structure of the Nucleosome Core Particle at 1.9Å Resolution††We dedicate this paper to the memory of Max Perutz who was particularly inspirational and supportive to T.J.R. in the early stages of this study. Journal of Molecular Biology. 319(5):1097–1113, doi: 10.1016/S0022-2836(02)00386-8, https://www.sciencedirect.com/science/article/pii/S0022283602003868.

32. Pettersen, E. F., T. D. Goddard, C. C. Huang, G. S. Couch, D. M. Greenblatt, E. C. Meng, and T. E. Ferrin. 2004. UCSF Chimera—A visualization system for exploratory research and analysis. Journal of Computational Chemistry. 25(13):1605–1612, doi: 10.1002/jcc.20084, https://onlinelibrary.wiley.com/doi/abs/10.1002/jcc.20084.

33. Ponder, J. W., and D. A. Case. 2003. Force fields for protein simulations. In Advances in Protein Chemistry, Protein Simulations. V. Daggett, editor. pp. 27-+.

34. Zgarbová, M., J. Sponer, M. Otyepka, T. E. Cheatham, R. Galindo-Murillo, and P. Jurecka. 2015. Refinement of the Sugar-Phosphate Backbone Torsion Beta for AMBER Force Fields Improves the Description of Z- and B-DNA. Journal of Chemical Theory and Computation. 11(12):5723–5736, doi: 10.1021/acs.jctc.5b00716, <GO to ISI>://WOS:000366223400016.

35. Tian, C., K. Kasavajhala, K. A. A. Belfon, L. Raguette, H. Huang, A. N. Migues, J. Bickel, Y. Z. Wang, J. Pincay, Q. Wu, and C. Simmerling. 2020. ff19SB: Amino-Acid-Specific Protein Backbone Parameters Trained against Quantum Mechanics Energy Surfaces in Solution. Journal of Chemical Theory and Computation. 16(1):528–552, doi: 10.1021/acs.jctc.9b00591, <GO to ISI>://WOS:000508474800041.

36. Izadi, S., R. Anandakrishnan, and A. V. Onufriev. 2014. Building Water Models: A Different Approach. Journal of Physical Chemistry Letters. 5(21):3863–3871, doi: 10.1021/jz501780a, <GO to ISI>://WOS:000344579500018.

37. Joung, I. S., and T. E. Cheatham. 2008. Determination of alkali and halide monovalent ion parameters for use in explicitly solvated biomolecular simulations. Journal of Physical Chemistry B. 112(30):9020–9041, doi: 10.1021/jp8001614, <GO to ISI>://WOS:000257926800026.

38. Kulkarni, M., C. Yang, and Y. Pak. 2018. Refined Alkali Metal Ion Parameters for the OPC Water Model. Bulletin of the Korean Chemical Society. 39(8):931–935, doi: 10.1002/bkcs.11527, <GO to ISI>://WOS:000440818400004.

39. Li, Z., L. F. Song, P. F. Li, and K. M. Merz. 2020. Systematic Parametrization of Divalent Metal Ions for the OPC3, OPC, TIP3P-FB, and TIP4P-FB Water Models. Journal of Chemical Theory and Computation. 16(7):4429–4442, doi: 10.1021/acs.jctc.0c00194, <GO to ISI>://WOS:000607532300037.

40. Case, D. A., T. E. Cheatham, T. Darden, H. Gohlke, R. Luo, K. M. Merz, A. Onufriev, C. Simmerling, B. Wang, and R. J. Woods. 2005. The Amber biomolecular simulation programs. Journal of Computational Chemistry. 26(16):1668–1688, doi: 10.1002/jcc.20290, <GO to ISI>://WOS:000233021400002.

41. Li, D. Z., Z. F. Chen, Z. J. Zhang, and J. Liu. 2017. Understanding Molecular Dynamics with Stochastic Processes via Real or Virtual Dynamics. Chinese Journal of Chemical Physics. 30(6):735–760, doi: 10.1063/1674-0068/30/cjcp1711223, <GO to ISI>://WOS:000423287900019.

42. Zhang, Z., X. Liu, K. Yan, M. E. Tuckerman, and J. Liu. 2019. Unified Efficient Thermostat Scheme for the Canonical Ensemble with Holonomic or Isokinetic Constraints via Molecular Dynamics. The Journal of Physical Chemistry A. 123(28):6056–6079, doi: 10.1021/acs.jpca.9b02771, 10.1021/acs.jpca.9b02771.

43. Berendsen, H. J. C., J. P. M. Postma, W. F. Van Gunsteren, A. Dinola, and J. R. Haak. 1984. Molecular dynamics with coupling to an external bath. The Journal of Chemical Physics. 81(8):3684–3690, doi: 10.1063/1.448118, 10.1063/1.448118.

44. Loncharich, R. J., B. R. Brooks, and R. W. Pastor. 1992. LANGEVIN DYNAMICS OF PEPTIDES - THE FRICTIONAL DEPENDENCE OF ISOMERIZATION RATES OF N-ACETYLALANYL-N’-METHYLAMIDE. Biopolymers. 32(5):523–535, doi: 10.1002/bip.360320508, <GO to ISI>://WOS:A1992HQ08500007.

45. Ryckaert, J.-P., G. Ciccotti, and H. J. C. Berendsen. 1977. Numerical integration of the cartesian equations of motion of a system with constraints: molecular dynamics of n-alkanes. Journal of Computational Physics. 23(3):327–341, doi: 10.1016/0021-9991(77)90098-5, 10.1016/0021-9991(77)90098-5.

46. Darden, T., D. York, and L. Pedersen. 1993. Particle mesh Ewald: An *N*⋅log(*N*) method for Ewald sums in large systems. The Journal of Chemical Physics. 98(12):10089–10092, doi: 10.1063/1.464397, 10.1063/1.464397.

47. Cornell, W. D., P. Cieplak, C. I. Bayly, I. R. Gould, K. M. Merz, D. M. Ferguson, D. C. Spellmeyer, T. Fox, J. W. Caldwell, and P. A. Kollman. 1995. A Second Generation Force Field for the Simulation of Proteins, Nucleic Acids, and Organic Molecules. Journal of the American Chemical Society. 117(19):5179–5197, doi: 10.1021/ja00124a002, 10.1021/ja00124a002.

48. Shaw, D. E., J. P. Grossman, J. A. Bank, B. Batson, J. A. Butts, J. C. Chao, M. M. Deneroff, R. O. Dror, A. Even, C. H. Fenton, A. Forte, J. Gagliardo, G. Gill, B. Greskamp, C. R. Ho, D. J. Ierardi, L. Iserovich, J. S. Kuskin, R. H. Larson, T. Layman, L.-S. Lee, A. K. Lerer, C. Li, D. Killebrew, K. M. Mackenzie, S. Y.-H. Mok, M. A. Moraes, R. Mueller, L. J. Nociolo, J. L. Peticolas, T. Quan, D. Ramot, J. K. Salmon, D. P. Scarpazza, U. B. Schafer, N. Siddique, C. W. Snyder, J. Spengler, P. T. P. Tang, M. Theobald, H. Toma, B. Towles, B. Vitale, S. C. Wang, and C. Young Anton 2: Raising the Bar for Performance and Programmability in a Special-Purpose Molecular Dynamics Supercomputer. IEEE.

49. Lippert, R. A., C. Predescu, D. J. Ierardi, K. M. Mackenzie, M. P. Eastwood, R. O. Dror, and D. E. Shaw. 2013. Accurate and efficient integration for molecular dynamics simulations at constant temperature and pressure. Journal of Chemical Physics. 139(16), 164106, doi: 10.1063/1.4825247, <GO to ISI>://WOS:000326637500009.

50. Martyna, G. J., D. J. Tobias, and M. L. Klein. 1994. CONSTANT-PRESSURE MOLECULAR-DYNAMICS ALGORITHMS. Journal of Chemical Physics. 101(5):4177–4189, doi: 10.1063/1.467468, <GO to ISI>://WOS:A1994PE11000084.

51. Evans, D. J., and B. L. Holian. 1985. THE NOSE-HOOVER THERMOSTAT. Journal of Chemical Physics. 83(8):4069–4074, doi: 10.1063/1.449071, <GO to ISI>://WOS:A1985ASA8600045.

52. Prescott, N. A., A. Mansisidor, Y. Bram, A. A. Lemmon, C. Lim, V. I. Risca, R. E. Schwartz, and Y. David (2023). A nucleosome switch primes Hepatitis B Virus infection. Cold Spring Harbor Laboratory.

53. Dyer, P. N., R. S. Edayathumangalam, C. L. White, Y. H. Bao, S. Chakravarthy, U. M. Muthurajan, and K. Luger. 2004. Reconstitution of nucleosome core particles from recombinant histones and DNA. In Methods in Enzymology, Chromatin and Chromatin Remodeling Enzymes, Pt A. C. D. Allis, and C. Wu, editors. pp. 23–44.

54. Taguchi, H., N. Horikoshi, Y. Arimura, and H. Kurumizaka. 2014. A method for evaluating nucleosome stability with a protein-binding fluorescent dye. Methods. 70(2-3):119–126, doi: 10.1016/j.ymeth.2014.08.019, <GO to ISI>://WOS:000346891000006.

55. Arimura, Y., K. Shirayama, N. Horikoshi, R. Fujita, H. Taguchi, W. Kagawa, T. Fukagawa, G. Almouzni, and H. Kurumizaka. 2014. Crystal structure and stable property of the cancer-associated heterotypic nucleosome containing CENP-A and H3.3. Scientific Reports. 4, 7115, doi: 10.1038/srep07115, <GO to ISI>://WOS:000346175800001.

56. Instruments, T. 2024. 16 November 2024 https://www.tainstruments.com/nanodsc/?gad_source=1&gclid=Cj0KCQiAouG5BhDBARIsAOc08RTmmlciQhbknpPBe2MbhIQBJ3VVaiFBtw1JaPsSOkj4C_dt3Fvcqw4aArqIEALw_wcB

57. Onufriev, A., D. Bashford, and D. A. Case. 2000. Modification of the generalized Born model suitable for macromolecules. Journal of Physical Chemistry B. 104(15):3712–3720, doi: 10.1021/jp994072s, <GO to ISI>://WOS:000086655600048.

58. Onufriev, A., D. Bashford, and D. A. Case. 2004. Exploring protein native states and large-scale conformational changes with a modified generalized born model. Proteins-Structure Function and Bioinformatics. 55(2):383–394, doi: 10.1002/prot.20033, <GO to ISI>://WOS:000220980600016.

59. Kollman, P. A., I. Massova, C. Reyes, B. Kuhn, S. Huo, L. Chong, M. Lee, T. Lee, Y. Duan, W. Wang, O. Donini, P. Cieplak, J. Srinivasan, D. A. Case, and T. E. Cheatham. 2000. Calculating Structures and Free Energies of Complex Molecules: Combining Molecular Mechanics and Continuum Models. Accounts of Chemical Research. 33(12):889–897, doi: 10.1021/ar000033j, 10.1021/ar000033j.

60. Srinivasan, J., T. E. Cheatham, P. Cieplak, P. A. Kollman, and D. A. Case. 1998. Continuum Solvent Studies of the Stability of DNA, RNA, and Phosphoramidate-DNA Helices. Journal of the American Chemical Society. 120(37):9401–9408, doi: 10.1021/ja981844, 10.1021/ja981844.

61. Miller, B. R., T. D. McGee, J. M. Swails, N. Homeyer, H. Gohlke, and A. E. Roitberg. 2012. *MMPBSA.py*: An Efficient Program for End-State Free Energy Calculations. Journal of Chemical Theory and Computation. 8(9):3314–3321, doi: 10.1021/ct300418h, <GO to ISI>://WOS:000308830700034.

62. Araya-Secchi, R., T. Perez-Acle, S. G. Kang, T. Huynh, A. Bernardin, Y. Escalona, J. A. Garate, A. D. Martínez, I. E. García, J. C. Sáez, and R. H. Zhou. 2014. Characterization of a Novel Water Pocket Inside the Human Cx26 Hemichannel Structure. Biophysical Journal. 107(3):599–612, doi: 10.1016/j.bpj.2014.05.037, <GO to ISI>://WOS:000340018000010.

63. Gowers, R. J., and P. Carbone. 2015. A multiscale approach to model hydrogen bonding: The case of polyamide. Journal of Chemical Physics. 142(22), 224907, doi: 10.1063/1.4922445, <GO to ISI>://WOS:000356176600039.

64. Gu, R. X., S. Baoukina, and D. P. Tieleman. 2019. Cholesterol Flip-Flop in Heterogeneous Membranes. Journal of Chemical Theory and Computation. 15(3):2064–2070, doi: 10.1021/acs.jctc.8b00933, <GO to ISI>://WOS:000461533000049.

65. Gowers, R., M. Linke, J. Barnoud, T. Reddy, M. Melo, S. Seyler, J. Domański, D. Dotson, S. Buchoux, I. Kenney, and O. Beckstein MDAnalysis: A Python Package for the Rapid Analysis of Molecular Dynamics Simulations. SciPy.

66. Michaud-Agrawal, N., E. J. Denning, T. B. Woolf, and O. Beckstein. 2011. MDAnalysis: A toolkit for the analysis of molecular dynamics simulations. Journal of Computational Chemistry. 32(10):2319–2327, doi: 10.1002/jcc.21787, 10.1002/jcc.21787.

67. Virtanen, P., R. Gommers, T. E. Oliphant, M. Haberland, T. Reddy, D. Cournapeau, E. Burovski, P. Peterson, W. Weckesser, J. Bright, S. J. Van Der Walt, M. Brett, J. Wilson, K. J. Millman, N. Mayorov, A. R. J. Nelson, E. Jones, R. Kern, E. Larson, C. J. Carey, İ. Polat, Y. Feng, E. W. Moore, J. Vanderplas, D. Laxalde, J. Perktold, R. Cimrman, I. Henriksen, E. A. Quintero, C. R. Harris, A. M. Archibald, A. H. Ribeiro, F. Pedregosa, P. Van Mulbregt, A. Vijaykumar, A. P. Bardelli, A. Rothberg, A. Hilboll, A. Kloeckner, A. Scopatz, A. Lee, A. Rokem, C. N. Woods, C. Fulton, C. Masson, C. Häggström, C. Fitzgerald, D. A. Nicholson, D. R. Hagen, D. V. Pasechnik, E. Olivetti, E. Martin, E. Wieser, F. Silva, F. Lenders, F. Wilhelm, G. Young, G. A. Price, G.-L. Ingold, G. E. Allen, G. R. Lee, H. Audren, I. Probst, J. P. Dietrich, J. Silterra, J. T. Webber, J. Slavič, J. Nothman, J. Buchner, J. Kulick, J. L. Schönberger, J. V. De Miranda Cardoso, J. Reimer, J. Harrington, J. L. C. Rodríguez, J. Nunez-Iglesias, J. Kuczynski, K. Tritz, M. Thoma, M. Newville, M. Kümmerer, M. Bolingbroke, M. Tartre, M. Pak, N. J. Smith, N. Nowaczyk, N. Shebanov, O. Pavlyk, P. A. Brodtkorb, P. Lee, R. T. McGibbon, R. Feldbauer, S. Lewis, S. Tygier, S. Sievert, S. Vigna, S. Peterson, S. More, T. Pudlik, T. Oshima, T. J. Pingel, T. P. Robitaille, T. Spura, T. R. Jones, T. Cera, T. Leslie, T. Zito, T. Krauss, U. Upadhyay, Y. O. Halchenko, and Y. Vázquez-Baeza. 2020. SciPy 1.0: fundamental algorithms for scientific computing in Python. Nature Methods. 17(3):261–272, doi: 10.1038/s41592-019-0686-2, 10.1038/s41592-019-0686-2.

68. Jolliffe, I. T., and J. Cadima. 2016. Principal component analysis: a review and recent developments. Philosophical Transactions of the Royal Society a-Mathematical Physical and Engineering Sciences. 374(2065), 20150202, doi: 10.1098/rsta.2015.0202, <GO to ISI>://WOS:000372553500008.

69. Amadei, A., M. A. Ceruso, and A. Di Nola. 1999. On the convergence of the conformational coordinates basis set obtained by the essential dynamics analysis of proteins’ molecular dynamics simulations. Proteins-Structure Function and Genetics. 36(4):419–424, doi: 10.1002/(sici)1097-0134(19990901)36:4<419::Aid-prot5>3.3.Co;2-l, <GO to ISI>://WOS:000081968000005 https://onlinelibrary.wiley.com/doi/10.1002/(SICI)1097-0134(19990901)36:4%3C419::AID-PROT5%3E3.0.CO;2-U.

70. Leo-Macias, A., P. Lopez-Romero, D. Lupyan, D. Zerbino, and A. R. Ortiz. 2005. An analysis of core deformations in protein superfamilies. Biophysical Journal. 88(2):1291–1299, doi: 10.1529/biophysj.104.052449, <GO to ISI>://WOS:000226750800046.

71. Sittel, F., A. Jain, and G. Stock. 2014. Principal component analysis of molecular dynamics: On the use of Cartesian vs. internal coordinates. Journal of Chemical Physics. 141(1), 014111, doi: 10.1063/1.4885338, <GO to ISI>://WOS:000339622000011.

72. Bakan, A., L. M. Meireles, and I. Bahar. 2011. ProDy: protein dynamics inferred from theory and experiments. Bioinformatics. 27(11):1575–1577, doi: 10.1093/bioinformatics/btr168.

73. Humphrey, W., A. Dalke, and K. Schulten. 1996. VMD: Visual molecular dynamics. Journal of Molecular Graphics & Modelling. 14(1):33–38, doi: 10.1016/0263-7855(96)00018-5, <GO to ISI>://WOS:A1996UH51500005.

74. Bakan, A., L. M. Meireles, and I. Bahar. 2011. *ProDy*: Protein Dynamics Inferred from Theory and Experiments. Bioinformatics. 27(11):1575–1577, doi: 10.1093/bioinformatics/btr168, 10.1093/bioinformatics/btr168.

75. Bakan, A., A. Dutta, W. Mao, Y. Liu, C. Chennubhotla, T. R. Lezon, and I. Bahar. 2014. *Evol* and *ProDy* for bridging protein sequence evolution and structural dynamics. Bioinformatics. 30(18):2681–2683, doi: 10.1093/bioinformatics/btu336, 10.1093/bioinformatics/btu336.

76. Grant, B. J., A. P. C. Rodrigues, K. M. ElSawy, J. A. McCammon, and L. S. D. Caves. 2006. Bio3d: an R package for the comparative analysis of protein structures. Bioinformatics. 22(21):2695–2696, doi: 10.1093/bioinformatics/btl461, <GO to ISI>://WOS:000241629600019.

77. Van Wynsberghe, A. W., and Q. Cui. 2005. Comparison of Mode Analyses at Different Resolutions Applied to Nucleic Acid Systems. Biophysical Journal. 89(5):2939–2949, doi: 10.1529/biophysj.105.065664, 10.1529/biophysj.105.065664.

78. Hayward, S. 2023. A Retrospective on the Development of Methods for the Analysis of Protein Conformational Ensembles. The Protein Journal. 42(3):181–191, doi: 10.1007/s10930-023-10113-9, 10.1007/s10930-023-10113-9.

79. Bahar, I., T. R. Lezon, A. Bakan, and I. H. Shrivastava. 2010. Normal Mode Analysis of Biomolecular Structures: Functional Mechanisms of Membrane Proteins. Chemical Reviews. 110(3):1463–1497, doi: 10.1021/cr900095e, 10.1021/cr900095e.

80. Atilgan, A. R., S. R. Durell, R. L. Jernigan, M. C. Demirel, O. Keskin, and I. Bahar. 2001. Anisotropy of Fluctuation Dynamics of Proteins with an Elastic Network Model. Biophysical Journal. 80(1):505–515, doi: 10.1016/s0006-3495(01)76033-x, 10.1016/s0006-3495(01)76033-x.

81. Skjærven, L., X.-Q. Yao, G. Scarabelli, and B. J. Grant. 2014. Integrating protein structural dynamics and evolutionary analysis with Bio3D. BMC Bioinformatics. 15(1), doi: 10.1186/s12859-014-0399-6, 10.1186/s12859-014-0399-6.

82. Bennett, R. L., A. Bele, E. C. Small, C. M. Will, B. Nabet, J. A. Oyer, X. X. Huang, R. P. Ghosh, A. T. Grzybowski, T. Yu, Q. Zhang, A. Riva, T. P. Lele, G. C. Schatz, N. L. Kelleher, A. J. Ruthenburg, J. Liphardt, and J. D. Licht. 2019. A Mutation in Histone H2B Represents a New Class of Oncogenic Driver. Cancer Discovery. 9(10):1438–1451, doi: 10.1158/2159-8290.Cd-19-0393, <GO to ISI>://WOS:000489623800026.

83. Wan, Y. C. E., and K. M. Chan. 2021. Histone H2B Mutations in Cancer. Biomedicines. 9(6):694, doi: 10.3390/biomedicines9060694, 10.3390/biomedicines9060694.

84. Ramaswamy, A., I. Bahar, and I. Ioshikhes. 2005. Structural dynamics of nucleosome core particle: Comparison with nucleosomes containing histone variants. Proteins-Structure Function and Bioinformatics. 58(3):683–696, doi: 10.1002/prot.20357, <GO to ISI>://WOS:000226695900017.

85. Bowerman, S., R. J. Hickok, and J. Wereszczynski. 2019. Unique Dynamics in Asymmetric macroH2A–H2A Hybrid Nucleosomes Result in Increased Complex Stability. The Journal of Physical Chemistry B. 123(2):419–427, doi: 10.1021/acs.jpcb.8b10668, 10.1021/acs.jpcb.8b10668.

86. Espiritu, D., A. K. Gribkova, S. Gupta, A. K. Shaytan, and A. R. Panchenko. 2021. Molecular Mechanisms of Oncogenesis through the Lens of Nucleosomes and Histones. Journal of Physical Chemistry B. 125(16):3963–3976, doi: 10.1021/acs.jpcb.1c00694, <GO to ISI>://WOS:000647271100001.

87. Ye, J., X. Ai, E. E. Eugeni, L. Zhang, L. R. Carpenter, M. A. Jelinek, M. A. Freitas, and M. R. Parthun. 2005. Histone H4 Lysine 91 Acetylation. Molecular Cell. 18(1):123–130, doi: 10.1016/j.molcel.2005.02.031, 10.1016/j.molcel.2005.02.031.

88. Yan, Q., S. Dutt, R. Xu, K. Graves, P. Juszczynski, J. P. Manis, and M. A. Shipp. 2009. BBAP Monoubiquitylates Histone H4 at Lysine 91 and Selectively Modulates the DNA Damage Response. Molecular Cell. 36(1):110–120, doi: 10.1016/j.molcel.2009.08.019, 10.1016/j.molcel.2009.08.019.

89. ThermoFisher. 2024. https://www.thermofisher.com/order/catalog/product/S6650

90. Bina, M., J. M. Sturtevant, and A. Stein. 1980. Stability of DNA in nucleosomes. Proceedings of the National Academy of Sciences. 77(7):4044–4047.

91. Kolomijtseva, G. Y., A. Prusov, E. Kolomijtseva, and T. Smirnova. 2023. Melting calorimetry of rat liver nuclei in the presence of magnesium ions. Biophysics. 68(2):272–281.

92. Balbi, C., M. L. Abelmoschi, L. Gogioso, S. Parodi, P. Barboro, B. Cavazza, and E. Patrone. 1989. Structural domains and conformational changes in nuclear chromatin: a quantitative thermodynamic approach by differential scanning calorimetry. Biochemistry. 28(8):3220–3227.

93. Cavazza, B., G. Brizzolara, G. Lazzarini, E. Patrone, M. Piccardo, P. Barboro, S. Parodi, A. Pasini, and C. Balbi. 1991. Thermodynamics of condensation of nuclear chromatin. A differential scanning calorimetry study of the salt-dependent structural transitions. Biochemistry. 30(37):9060–9072.

94. Kang, T. Z. E., L. Zhu, D. Yang, D. Ding, X. Zhu, Y. C. E. Wan, J. Liu, S. Ramakrishnan, L. L. Chan, S. Y. Chan, X. Wang, H. Gan, J. Han, T. Ishibashi, Q. Li, and K. M. Chan. 2021. The elevated transcription of ADAM19 by the oncohistone H2BE76K contributes to oncogenic properties in breast cancer. Journal of Biological Chemistry. 296:100374, doi: 10.1016/j.jbc.2021.100374, 10.1016/j.jbc.2021.100374.

95. Kimura, T., S. Hirai, T. Kujirai, R. Fujita, M. Ogasawara, H. Ehara, S. i. Sekine, Y. Takizawa, and H. Kurumizaka. 2024. Cryo-EM structure and biochemical analyses of the nucleosome containing the cancer-associated histone H3 mutation E97K. Genes to Cells. 29(9):769–781.

96. Sun, R., Z. Li, and T. C. Bishop. 2019. TMB library of nucleosome simulations. Journal of chemical information and modeling. 59(10):4289–4299.

97. Das, C., and J. K. Tyler. 2012. Histone exchange and histone modifications during transcription and aging. Biochimica Et Biophysica Acta-Gene Regulatory Mechanisms. 1819(3-4):332–342, doi: 10.1016/j.bbagrm.2011.08.001, <GO to ISI>://WOS:000301628800015.

98. Cheung, V., G. Chua, N. N. Batada, C. R. Landry, S. W. Michnick, T. R. Hughes, and F. Winston. 2008. Chromatin- and Transcription-Related Factors Repress Transcription from within Coding Regions throughout the *Saccharomyces cerevisiae* Genome. Plos Biology. 6(11):2550–2562, e277, doi: 10.1371/journal.pbio.0060277, <GO to ISI>://WOS:000261187900021.

99. Jackson, V. 1988. Deposition of newly synthesized histones: hybrid nucleosomes are not tandemly arranged on daughter DNA strands. Biochemistry. 27(6):2109–2120, doi: 10.1021/bi00406a044.

100. Schwabish, M. A., and K. Struhl. 2004. Evidence for eviction and rapid deposition of histories upon transcriptional elongation by RNA polymerase II. Molecular and Cellular Biology. 24(23):10111–10117, doi: 10.1128/mcb.24.23.10111-10117.2004, <GO to ISI>://WOS:000225167800002.

101. Lee, C. K., Y. Shibata, B. Rao, B. D. Strahl, and J. D. Lieb. 2004. Evidence for nucleosome depletion at active regulatory regions genome-wide. Nature Genetics. 36(8):900–905, doi: 10.1038/ng1400, <GO to ISI>://WOS:000222974000026.

