## Supplementary Information for "Structural and Thermodynamic Impact of Oncogenic Mutations on the Nucleosome Core Particle"

**This PDF file includes:**

Supporting text
Figures SI to S15
Tables S1 to S2
SI References

**Other supporting materials for this manuscript include the following:**

Movies S1 to S3
WT, E76K, and R92T simulations, focusing on the H2B-H4 interface

**Supplementary Information Text**

**Analysis of MD Simulations at 2.4 M NaCl**

The alcohol group of H4T92 in the H4R92T system is too short and not polar enough to form a hydrogen bond with H2BE76, thereby decreasing the stability of the H4-H2B interface compared with the *WT* system. Also, Lehmann and co-workers showed that mutation of H2R88 substantially decreases nucleosome assembly, revealing that specific arginines in the histone core are vital for nucleosome stability (1). As the propensity for forming hydrogen bonds between amino acids in the H2B α2 and H4 α3 helices decreases, the frequency of forming hydrogen bonds at other helix interfaces in the H2B and H4 interface increases. Destroying the hydrogen bond between H4T92 and H2BE76 in the H4R92T system, promoted a hydrogen bond between H4Y72 and H2BE76 with a lifetime of 7.1 ps (**Figure 2a**). The probability of the alcohol group of H4T71 interacting with the guanidino group of H2BR99 also increased in the H2BE76K mutation.

At high salt concentration of 2.4 M, the probability of forming the salt bridge between H2BE76 and H4R92 was drastically reduced, however the hydrogen bond between H2BY83 and H4Y88 was sustained. The **π-π** interaction of the two tyrosines (H2BY83 and H4Y88) was preserved (Figure S1a-c). At 2.4 M the H2BE76K and H4R92T systems also showed a lower frequency of hydrogen bonding and salt bridge formation between interacting amino acids on the H2B-α2 and H4-α3 helices. Like the results at physiological condition, our mutations weakened interhelical hydrogen bonding and salt bridge formation at one position while promoting such interactions at other positions in the H4-H2B interface (Figure S2). The distances of the center of geometry of the backbones (alpha carbon) of the interacting amino acids (H2BE76-H4R92, H2BY83-H4K91) or the phenolic groups of the interacting tyrosine (H2BY83-H4Y88) were lower in the *WT* system than both mutants at 0.15 M (Figure S4). Distance-angle two-dimensional plot of the phenolic rings of H2BY83 and H4Y88 involved in **π-π** interactions showed that at 0.15 M, the rings maintained one orientation angle in the *WT* systems but sampled between two angles in the H2BE76K system (Figure S6a). This means that the rings are more stable in the *WT* system than in the H2BE76K system. Increasing the salt concentration of *NaCl* to 2.4 M affected the orientation of the rings (Figure S6b). The orientation of the phenolic rings for the H4R92T system display a peak at approximately 3° (Figure S6).

**Supplementary Table 1:** Molecular Dynamics Simulation Systems

| System Properties | WT_0.15M | WT_2.4M | H2BE76K_0.15M | H2BE76K_2.4M | H4R92T_0.15M | H4R92T_2.4M |
|---|---|---|---|---|---|---|
| **Box Dimension** | 167x201x117 | 147x155x135 | 167x201x117 | 159x191x112 | 167x201x117 | 167x201x117 |
| **Box Volume equilibrated (Å$^3$)** | 3941876.356 | 3081213.081 | 3941957.576 | 3317870.103 | 3941790.792 | 3941790.792 |
| **No. of Atoms** | 444888 | 396256 | 444911 | 425809 | 444799 | 412753 |
| **No. of Water** | 104740 | 90508 | 104745 | 97895 | 104720 | 94038 |
| **No. of Na+** | 472 | 4620 | 469 | 4618 | 473 | 5814 |
| **No. of Cl-** | 356 | 4504 | 355 | 4504 | 356 | 5697 |
| **No. of Mg2+** | 14 | 14 | 14 | 14 | 14 | 14 |
| **[NaCl]** | 0.15 | 2.4 | 0.15 | 2.4 | 0.15 | 2.4 |
| **Equilibration Time** | 100 ns | 100ns | 100ns | 100ns | 100ns | 100ns |
| **Equilibration done on** | AMBER | AMBER | AMBER | AMBER | AMBER | AMBER |
| **Production Run** | 6us | 12us | 6us | 12us | 6us | 6us |
| **Production run done on** | Anton2 | Anton2 | Anton2 | Anton2 | Anton2 | Anton2 |

**Supplementary Table 2:** Primer Sequence for Replicating Mutant Histones

| Systems | Forward Sequence | Reverse Sequence |
|---|---|---|
| H2BE76K | 5’-TCGCAGGGAAAGCCTCCCGCCTGGCTC-3’ | 5’-AGGCTTTCCCTGCGATGCGCTCAAAC-3’ |
| H4R92T | 5’-TCTGAAAACCCAGGGTCGTACCCTGTACGG-3’ | 5’-CCCTGGGTTTTCAGAGCGTAAACAACGTCC-3’ |

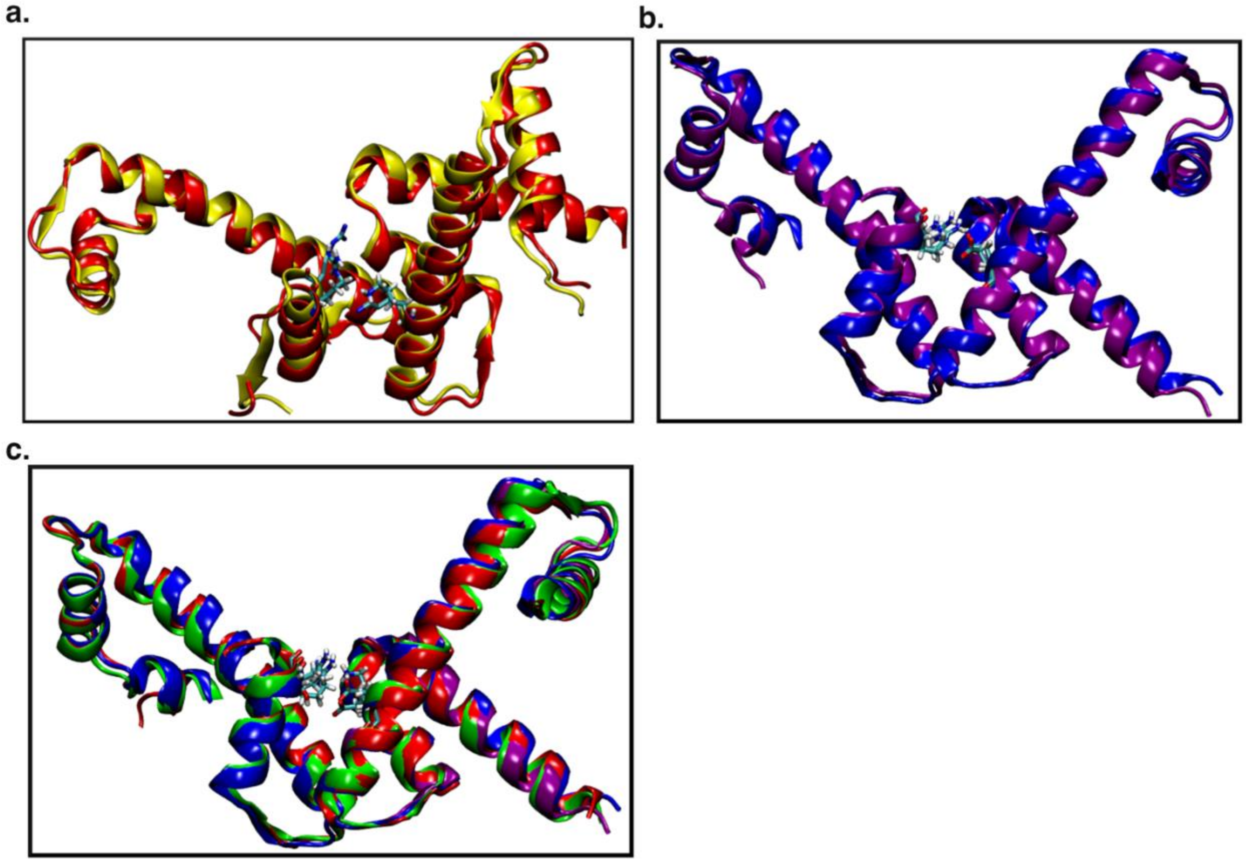

**Suppl. Figure 1:** (a) Overlayed structure of the H2BE76K simulated system (red) and the H2BE76K crystal structure (yellow). H2BK76 and H4R92 shown in licorice. (b) Overlayed structure of the WT simulated system (blue) and the 1KX5 crystal structure (in magenta). H2BE76 and H4R92 shown in licorice. (c) Histone H4-H2B alignment for all simulated systems (WT, H2BE76K and H4R92T) at 6$\boldsymbol{\mu}$s. The *WT* system is color blue, H2BE76K is in red and H4R92T is colored green. Amino acids H2BE76/H2BK76 and H4R92/H4T92 are shown in licorice.

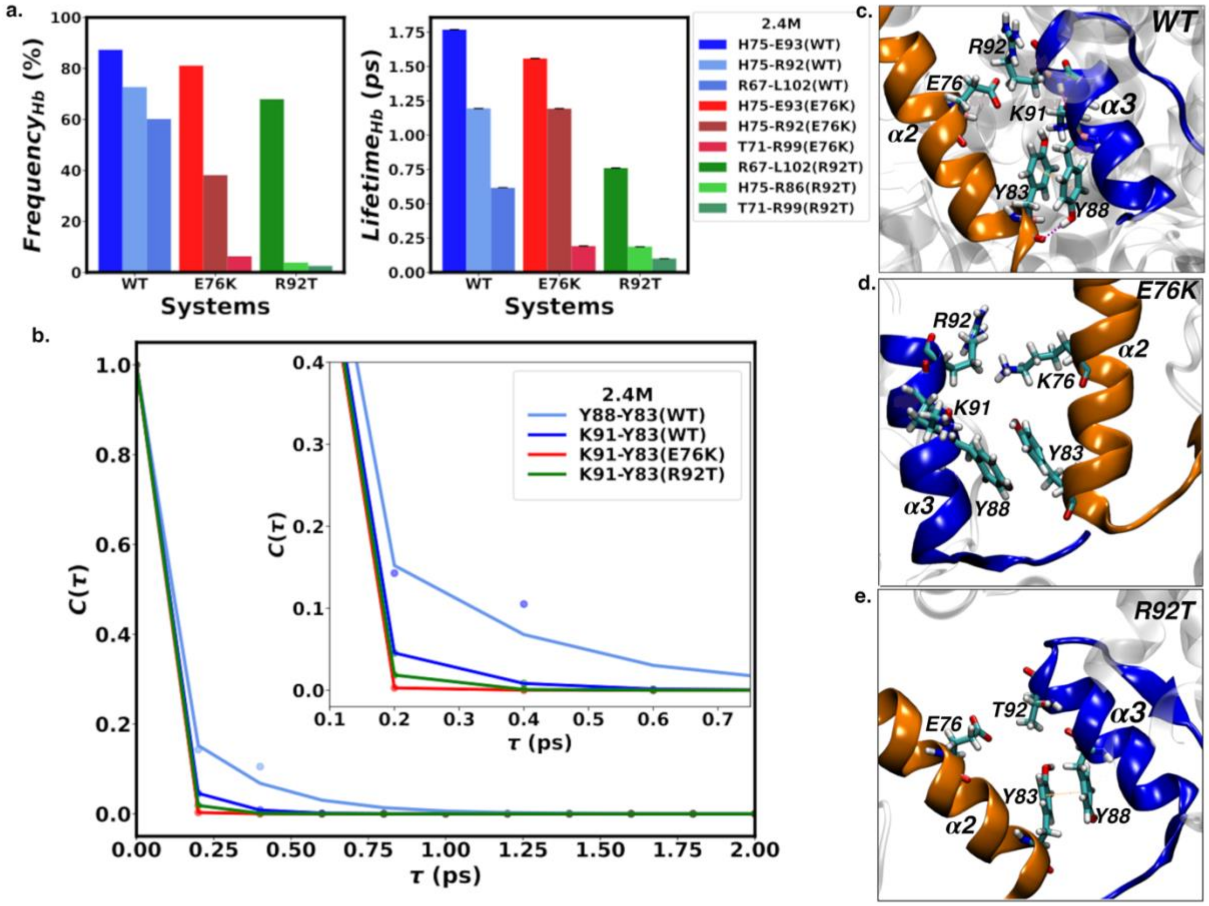


**Suppl. Figure 2:** (a) The left panel shows the percentage frequency of hydrogen bonds at the H4-H2B interface for the simulation, while the right panel shows the continuous lifetime of each hydrogen bond type. The amino acid key in "a" is written in the format: H4-H2B(system). (b) Time autocorrelation function for hydrogen bonds and salt bridges between **α**2 (H2B) and **α**3 (H4) helices at the H4-H2B interface at 2.4 M NaCl. The salt bridge between H2BE76 and H4R92 at this interface was weakened by an increase in the concentration of NaCl in the system. This salt bridge was also absent in the mutant systems (H2BE76K and H4R92T). The panel key shows amino acids labeled in H4-H2B(system) format. (c) Amino acid configuration in the *WT* system at 2.4 M showing the hydrogen bonds and **π-π** interaction between interacting amino acids on the orange **α**2 helix on H2B and the blue **α**3 helix on H4 (d) The **π-π** interaction and hydrogen bonds between interacting amino acids were broken in the H2BE76K system at 2.4 M (e) The **π-π** interaction is preserved in the H4R92T system, but H4T92 and H2BE76 do not take part in any direct electrostatic interaction.

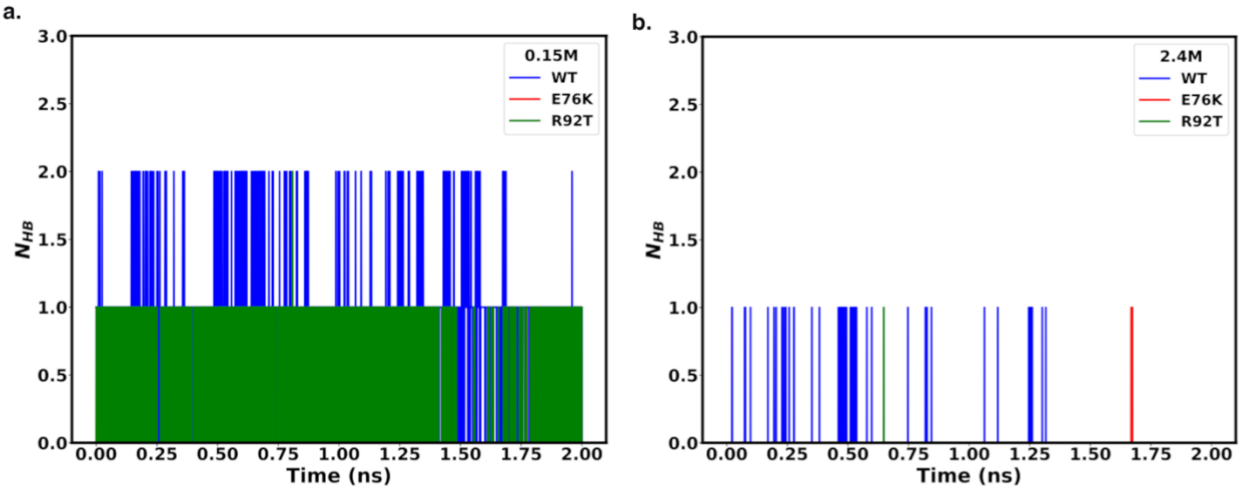


**Suppl. Figure 3:** The total number of hydrogen bonds or salt bridges between **α**2 (H2B) and **α**3 (H4) helices at the H4-H2B interface for all systems at (a) physiological condition of 0.15 M and (b) high salt concentration of 2.4 M. The total number of hydrogen bonds formed between the interacting amino acids in the **α**2 helix in H2B and **α**3 helix in H4 was higher for the wild-type system when compared to the mutant systems (H2BE76K and H4R92T). The H4R92T system had slightly more stable electrostatic interactions when compared with the E76K system.

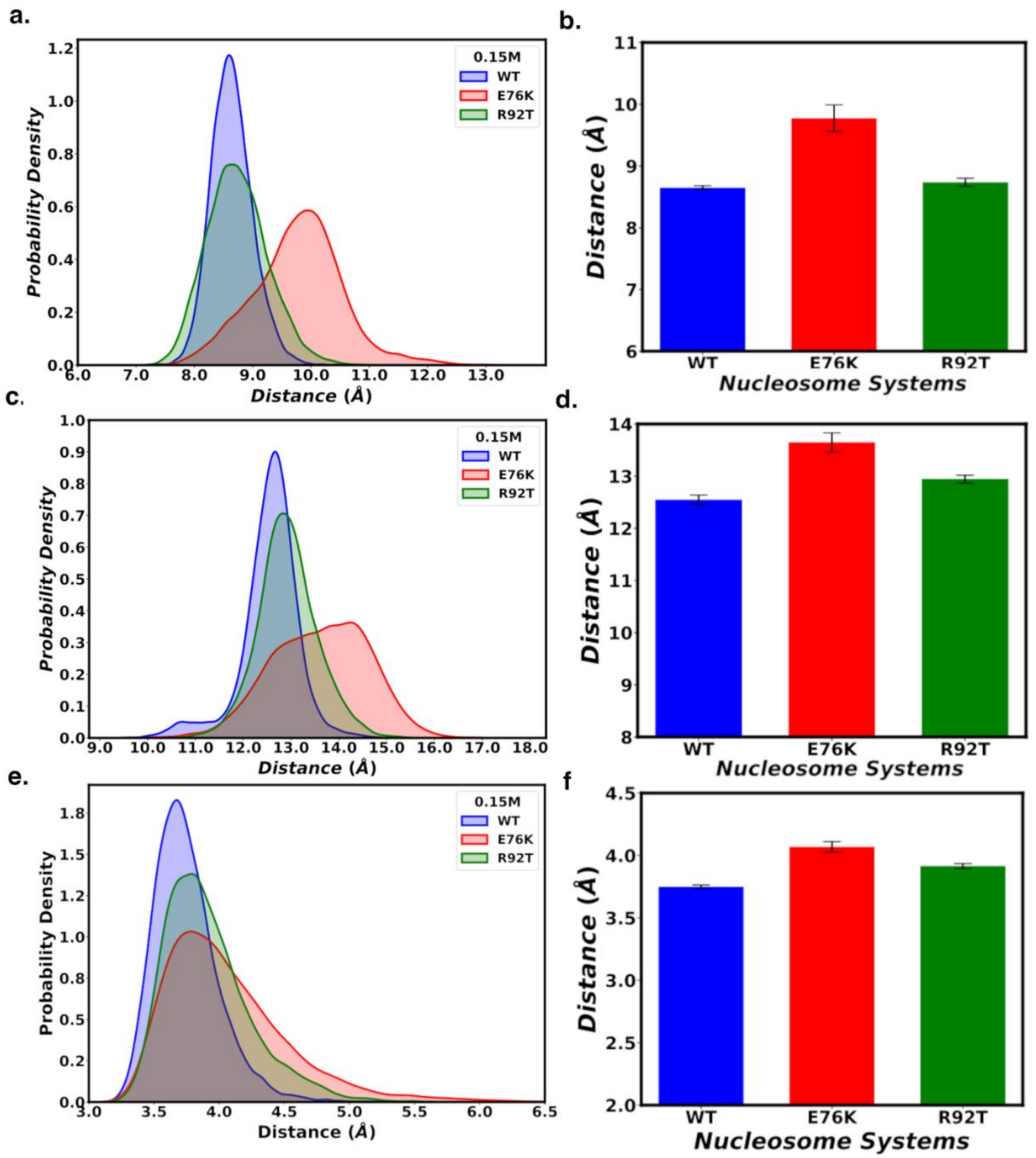


**Suppl. Figure 4:** (a) and (b) The distance between the alpha carbons of H2BE76 and H4R92 at 0.15 M for all systems. (c) and (d) The distance between the alpha carbons of H2BY83 and H4K91 at 0.15 M for all systems. (e) and (f) The distance between the plane of the aromatic rings of H2BY83 and H4Y88 at 0.15 M for all systems. The probability distributions revealed that the distance between the alpha carbons or aromatic rings of interacting amino acids at the **α**2 (H2B) and **α**3 (H4) helices in the H4-H2B interface is lower for the wild-type (*WT*) system compared to the H2BE76K mutant systems. The closer the interacting amino acids in the wild-type system, the higher the probability of forming hydrogen bonds, salt bridges, and **π-π** interactions.

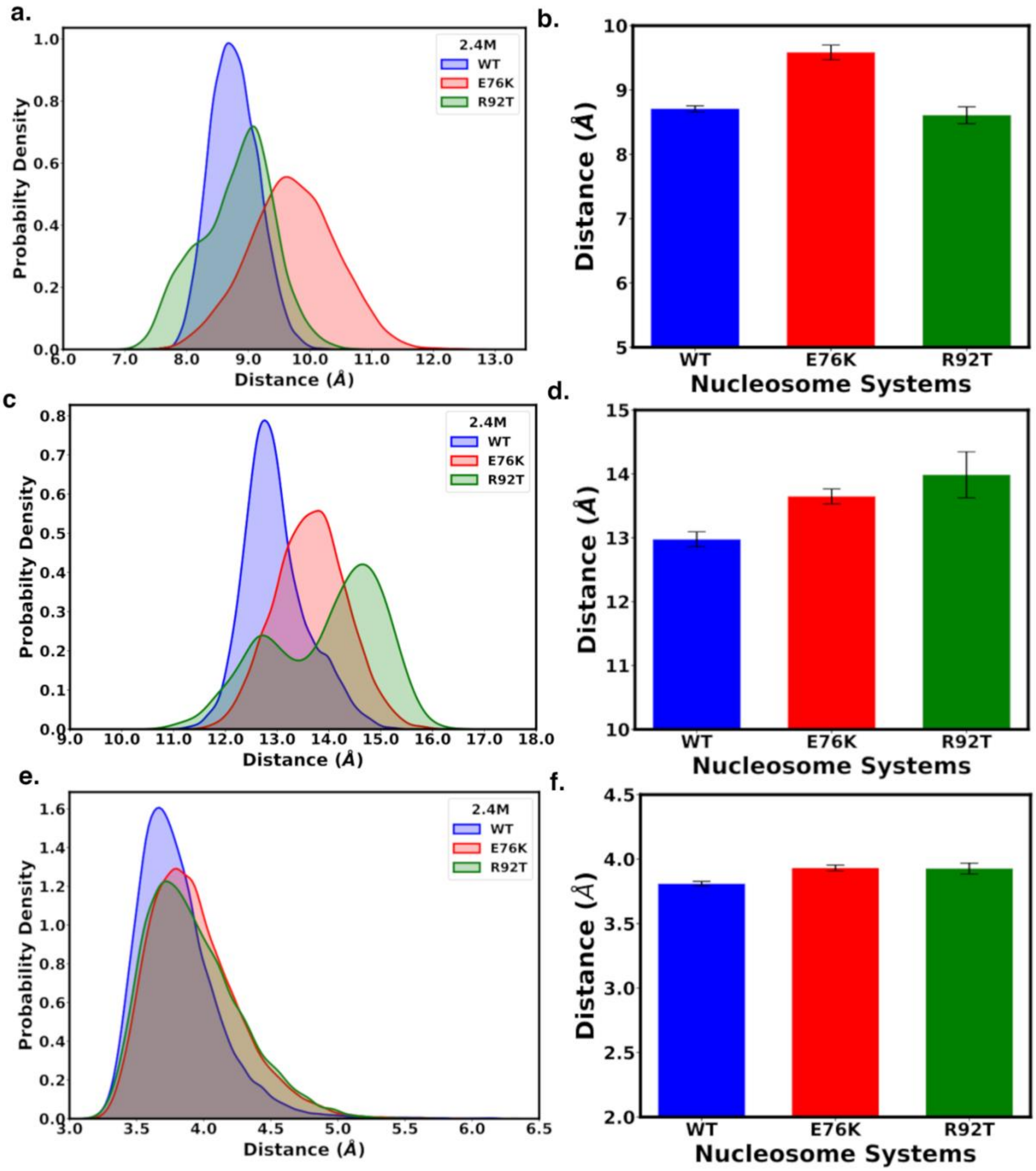


**Suppl. Figure 5:** (a) and (b) The distance between the alpha carbons of H2BE76 and H4R92 at 2.4 M for all systems. (c) and (d) The distance between the alpha carbons of H2BY83 and H4K91 at 2.4 M for all systems. (e) and (f) The distance between the plane of the aromatic rings of H2BY83 and H4Y88 at and 2.4 M for all systems. The probability distributions revealed that the distance between the alpha carbons or aromatic rings of interacting amino acids at the **α**2 (H2B) and **α**3 (H4) helices in the H4-H2B interface is lower for the wild-type (*WT*) system compared to the mutant systems (H2BE76K and H4R92T). The distance between H2BE76 and H4R92 in the WT and H4R92T system were somewhat similar.

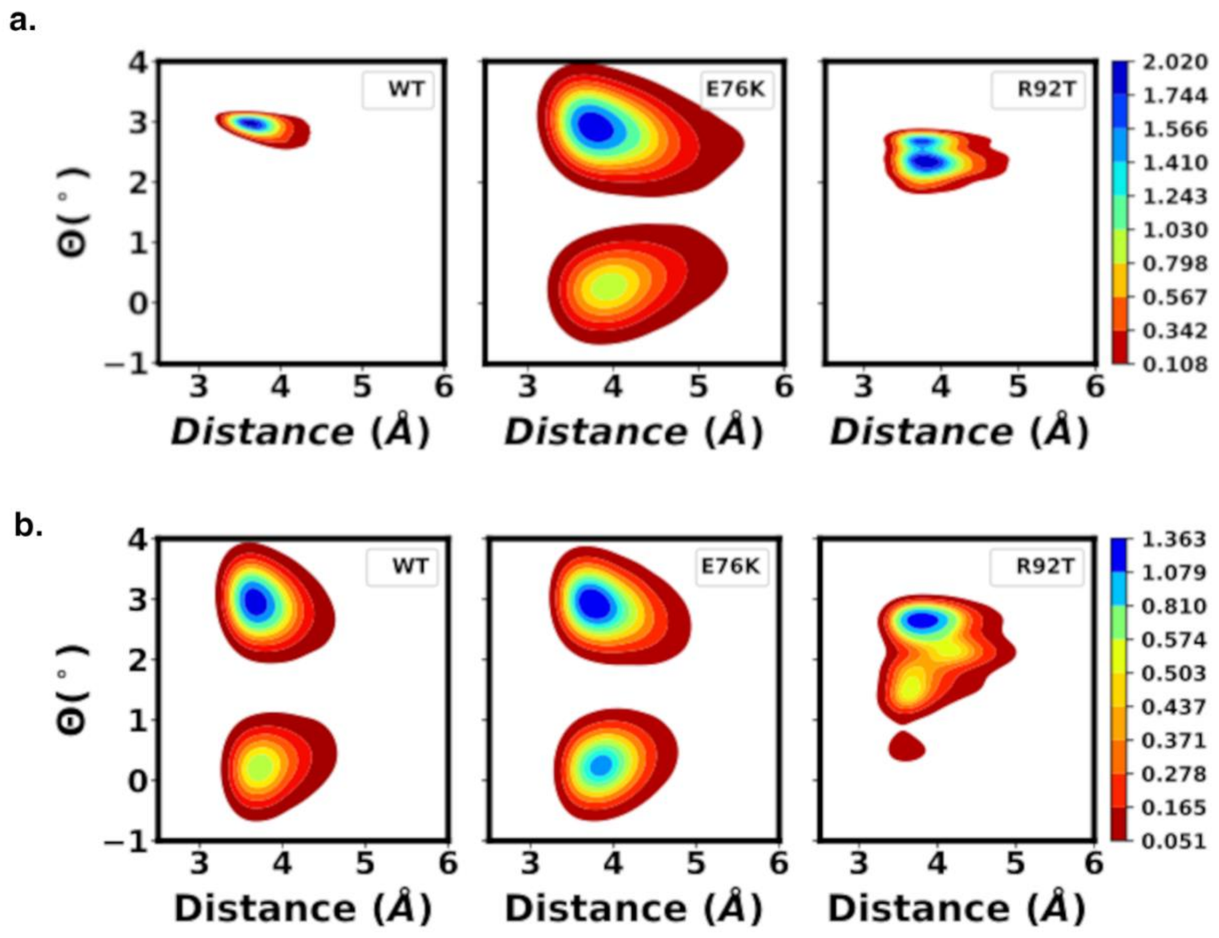


**Suppl. Figure 6:** (a) and (b) Angle-distance two-dimensional plot of the dihedral angle between the plane of the aromatic rings in H2BY83 and H4Y88 at 0.15 M and 2.4 M. The wild-type (*WT*) system maintained one conformational state at physiological conditions of 0.15 M NaCl, while the H2BE76K mutant system navigated between two distinct conformations. The aromatic rings in the H4R92T system also showed two somewhat similar configurations that were inter-convertible. At a high salt concentration of 2.4 M, all systems showed two conformations of the aromatic rings.

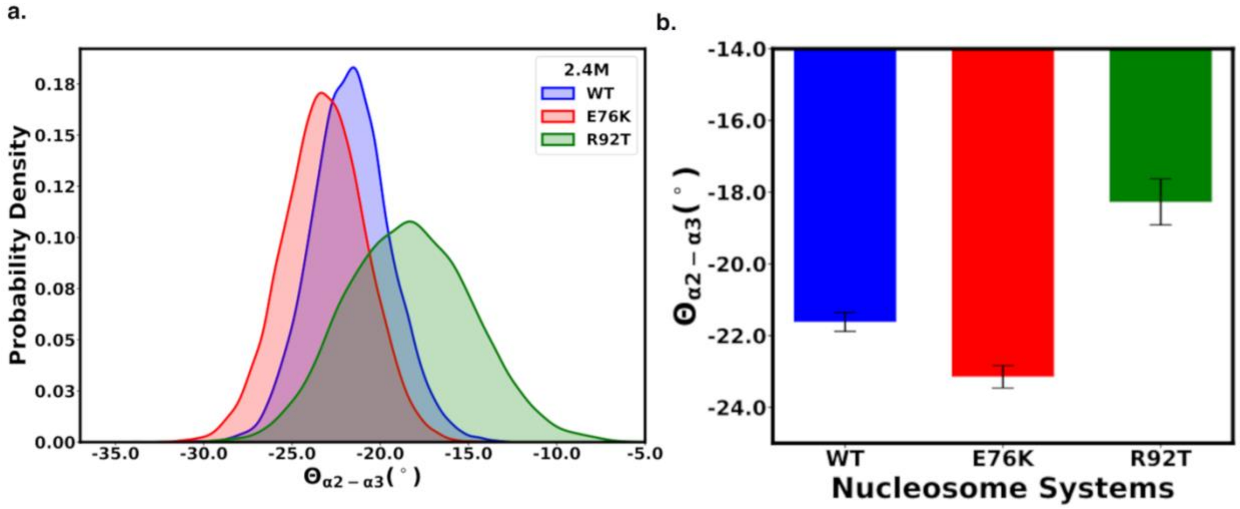


**Suppl. Figure 7:** (a) and (b) Dihedral angle in all systems at 2.4 M. The helix angle between H2B **α**2 and H4 **α**3 helices in the mutants (H2BE76K and H4R92T) were different from the *WT* system.

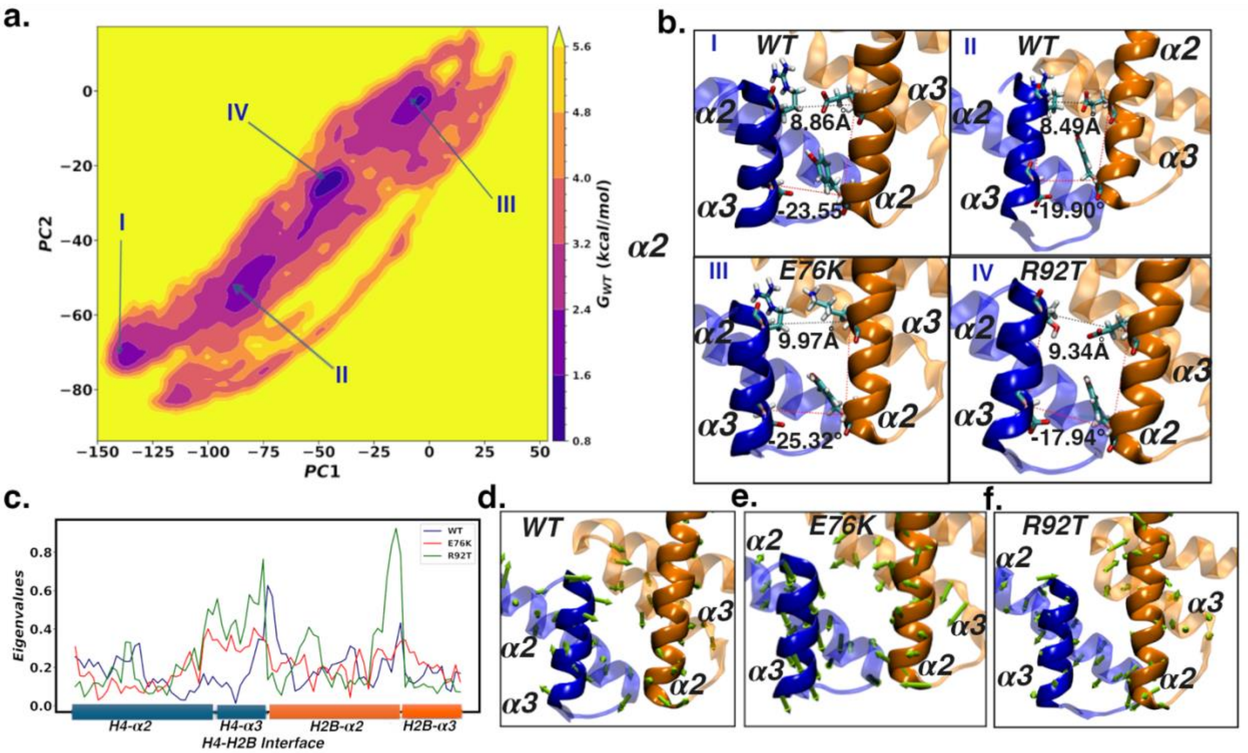


**Suppl. Figure 8:** Principal component analysis (PCA) for the combined systems using the coordinates of α2 and α3 helices of both H4 and H2B at the H4-H2B interface at 2.4 M of NaCl. The free-energy plot showed energy minima representing each system as I, II, III and IV. (b) The α2 and α3 helix conformations at the H4-H2B interface for WT, H2BE76K and H4R92T systems. The distance and dihedral angles between the plane of the helices are shown. At 2.4 M, the interhelical distance between the WT and H4R92T systems were statistically similar but there was a slight increase of approximately 0.3-0.5 Å in the H2BE76K system when compared to the WT system. The dihedral angle of the planes of the H2B-α2 and H4-α3 helices in the H4R92T system was different from the WT and H2BE76K system. (c) Eigenvalues at the H4-H3B interface showed H2B-α2 and H4-α3 in H2BE76K and H4R92T had the highest fluctuation. Eigenvector representation of the PC for (d) WT, (e) H2BE76K and (f) H4R92T systems. The amplitude of divergence of the helices in the H2BE76K system decreased at 2.4M when compared to physiological concentration of 0.15M of NaCl. There was still a correlation in the movement of the helices in the WT and H4R92T systems.

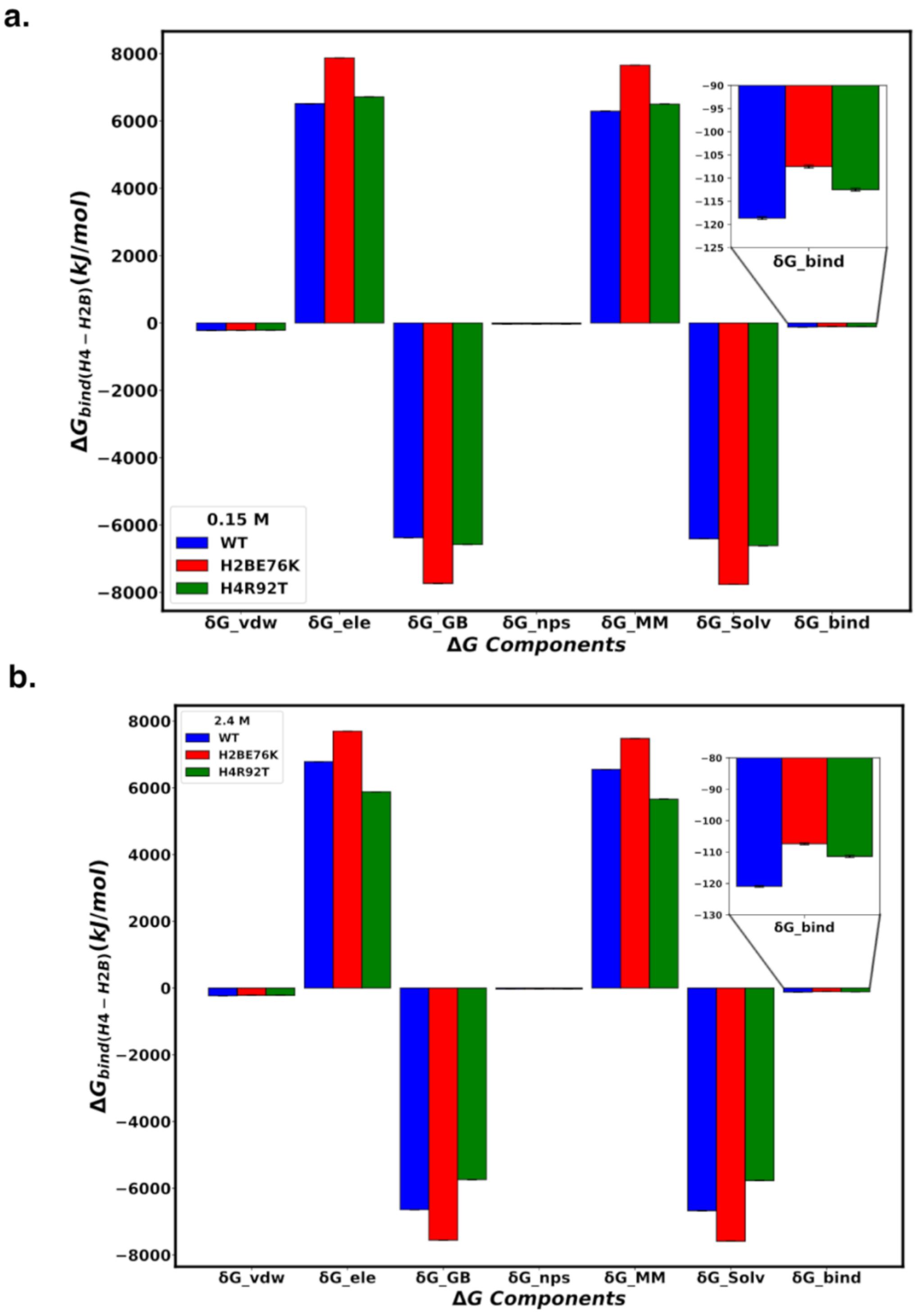


**Suppl. Figure 9:** (a) and (b) Contribution of different energy components to the binding free energy at 0.15 M and 2.4 M respectively. The binding free energy between histone H2B and H4 is driven by electrostatic contributions in the histones ($\Delta E_{elec}$) and the ionic solvation free energy component between the histones and the solvent ($\Delta G_{GB}$). The energy contribution of non-polar ($\Delta G_{nps}$) or van der Waal's ($\Delta E_{vdw}$) interactions is minimal.

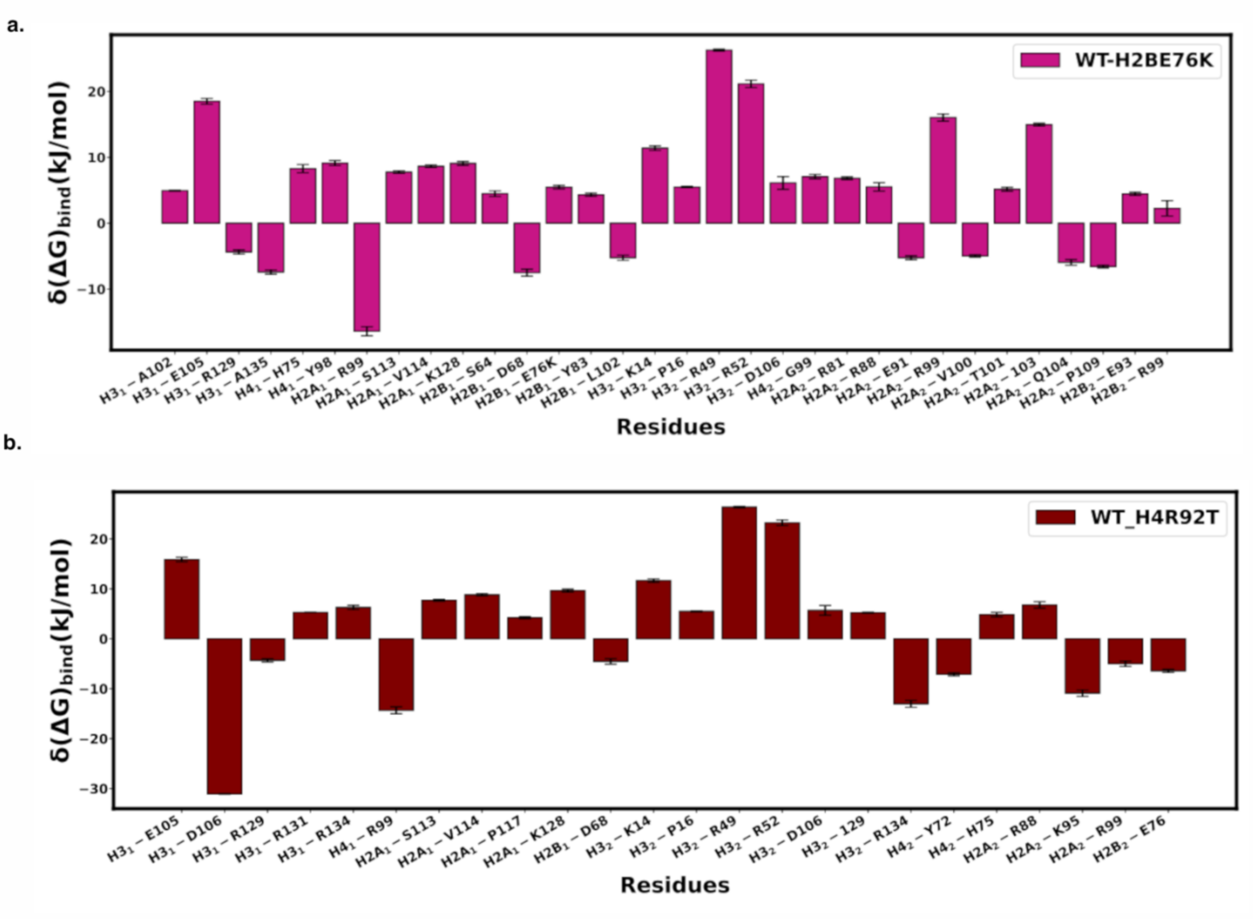


**Suppl. Figure 10:** Variation in the amino acid's contributions to the binding free energy at the Tetramer (H3-H4)-Dimer (H2A-H2B) interface upon mutation. Globally, mutation affected the amino acid contribution to the binding free energy at other interfaces beside the point of mutation.

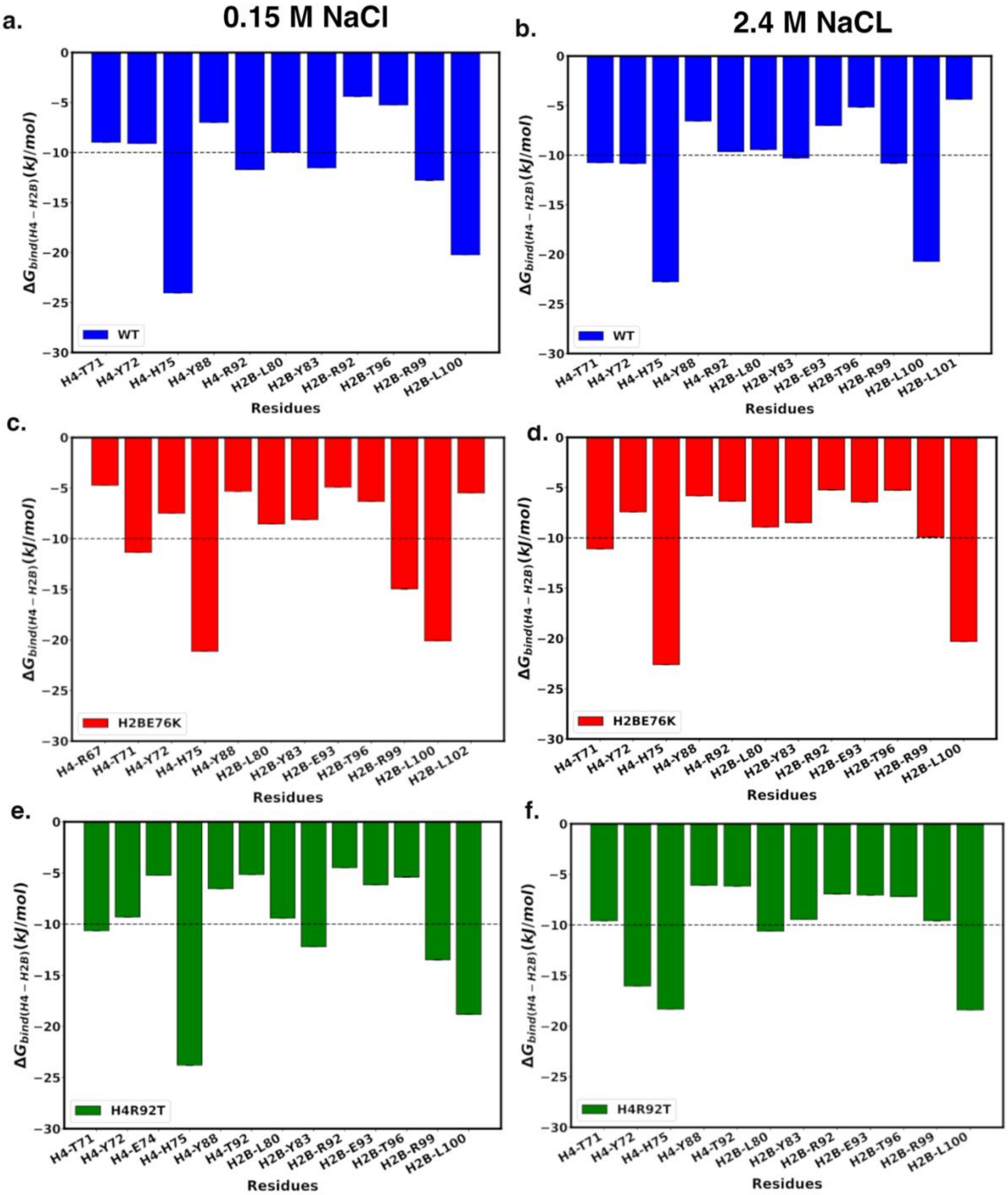


**Suppl. Figure 11:** Amino acid residue contribution to the binding free energy between histone H4 and H2B for (a) Wild-type system at 0.15 M (b) Wild-type system at 2.4 M (c) H2BE76K system at 0.15 M (d) H2BE76K system at 2.4 M (e) H4R92T system at 0.15 M (b) H4R92T system at 2.4 M. H4H75, H4R92, H2BR99 and H2BL100 were the main contributors to the binding free energy. Energy contributions H4Y88, H4R92, and H2BY83 were decreased by mutation. However, mutation improved the stability of H4T71 and H2BR99 in the H2BE76K system and H4Y72 and H2BR99in the H4R92T system.

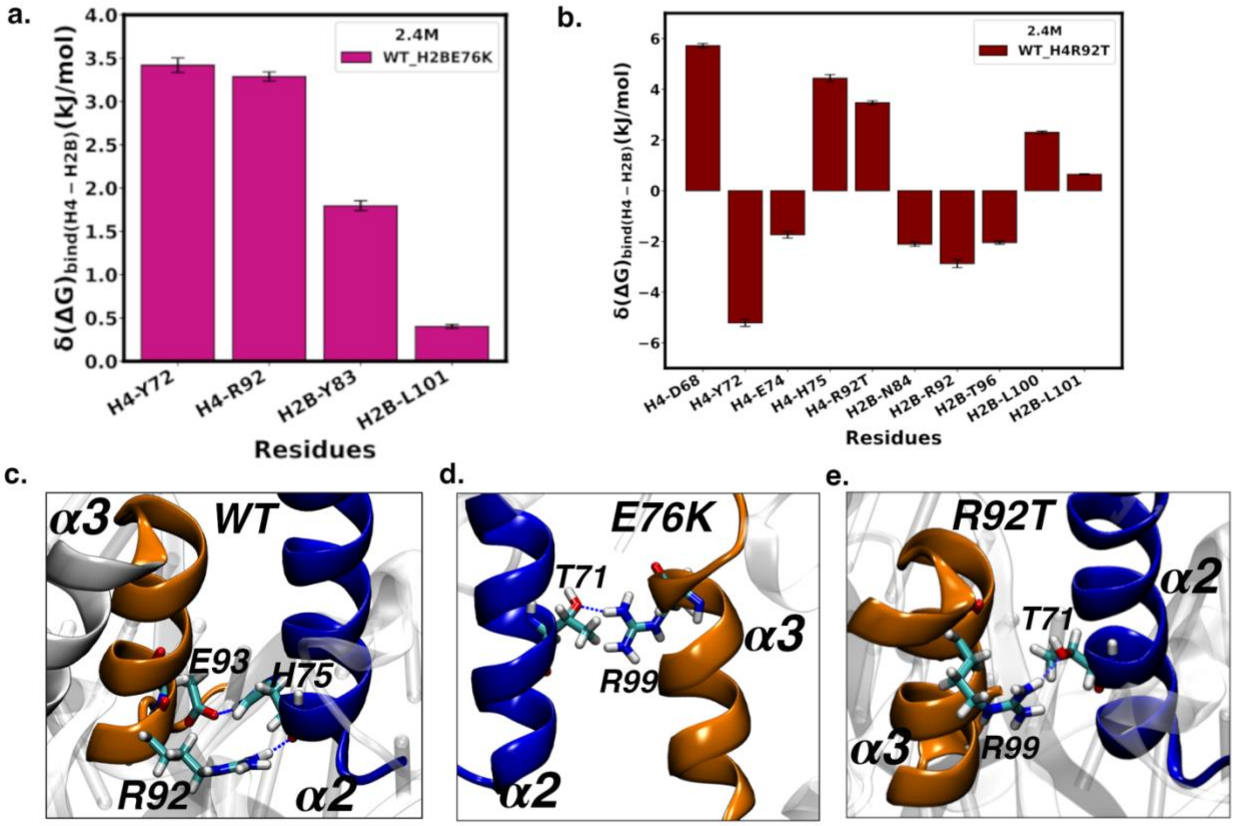


**Suppl. Figure 12:** (a) and (b) Variation in the decomposition free energy contribution between histone H4 and H2B upon mutation at 2.4 M NaCl. The amino acids most affected were at the **α**2 and **α**3 helices of H4 and H2B. The amino acid contribution to the binding free energy for H4R92, H2BE76 and H2BY83 decreased upon H2BE76K and H4R92T mutations. The contribution of H4Y72 improved in the H4R92T mutant but reduced in the H2BE76K mutant.

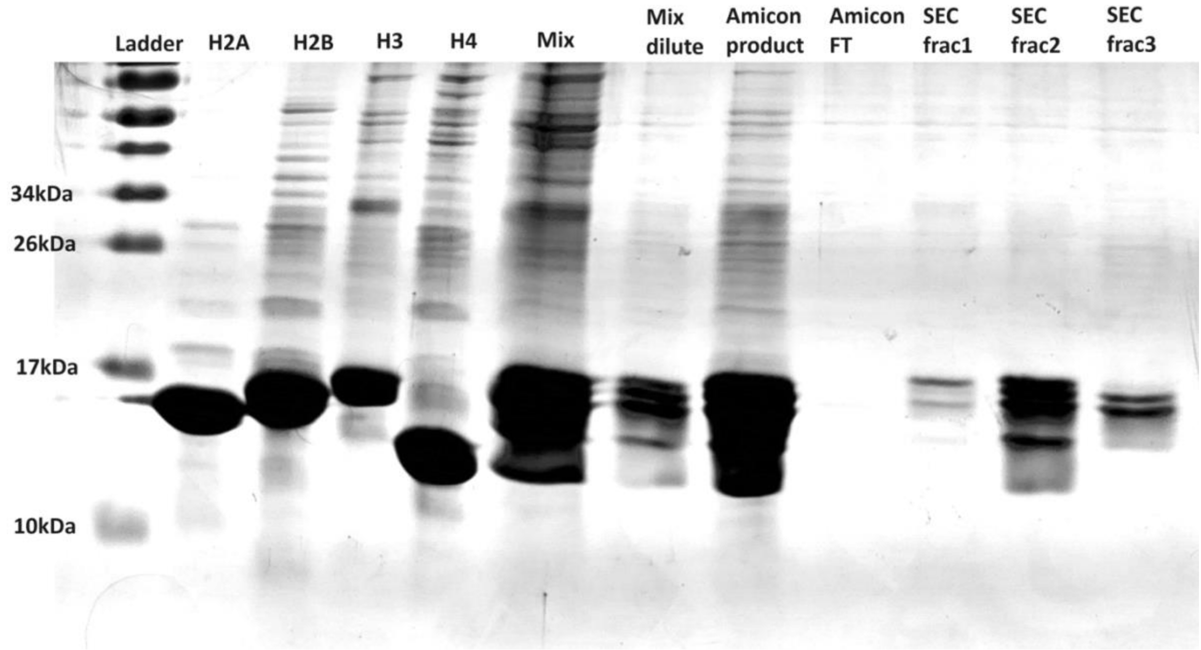


**Suppl. Figure 13:** SDS-PAGE gel of pure histone samples and WT octamer refolding. Lanes 2-5 are redissolved lyophilized pure histones purified by reverse-phase HPLC. Lanes 6-12 correspond to steps in the octamer refolding protocol where lane 11 (SEC frac2) is octamer purified by Size Exclusion FPLC.

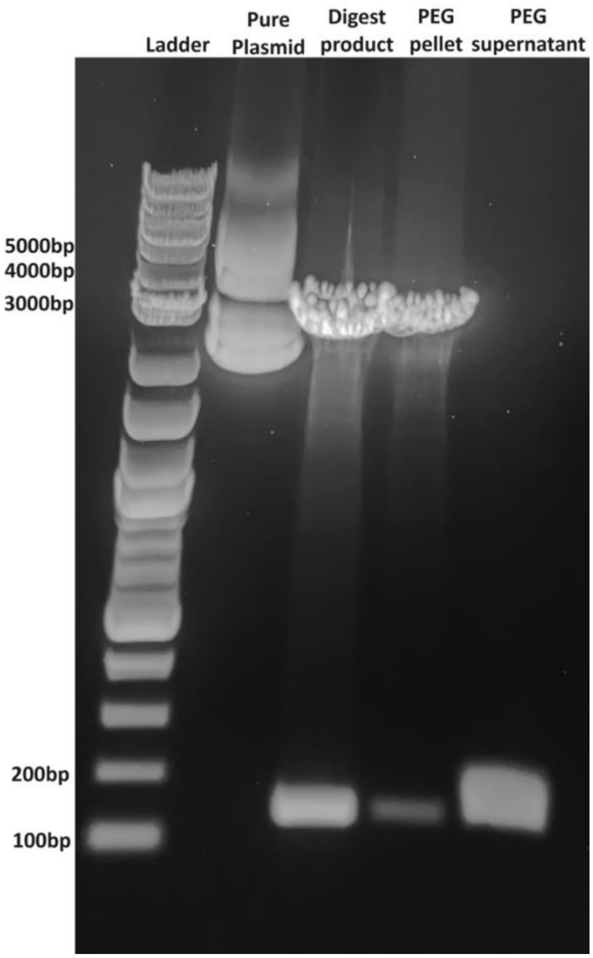


**Suppl. Figure 14:** 1% Acrylamide gel showing purification of 12 copy WIDOM 601 (147bp) plasmid and digestion product. The full intact plasmid is 4599 bp and the digested backbone is 2835 bp.

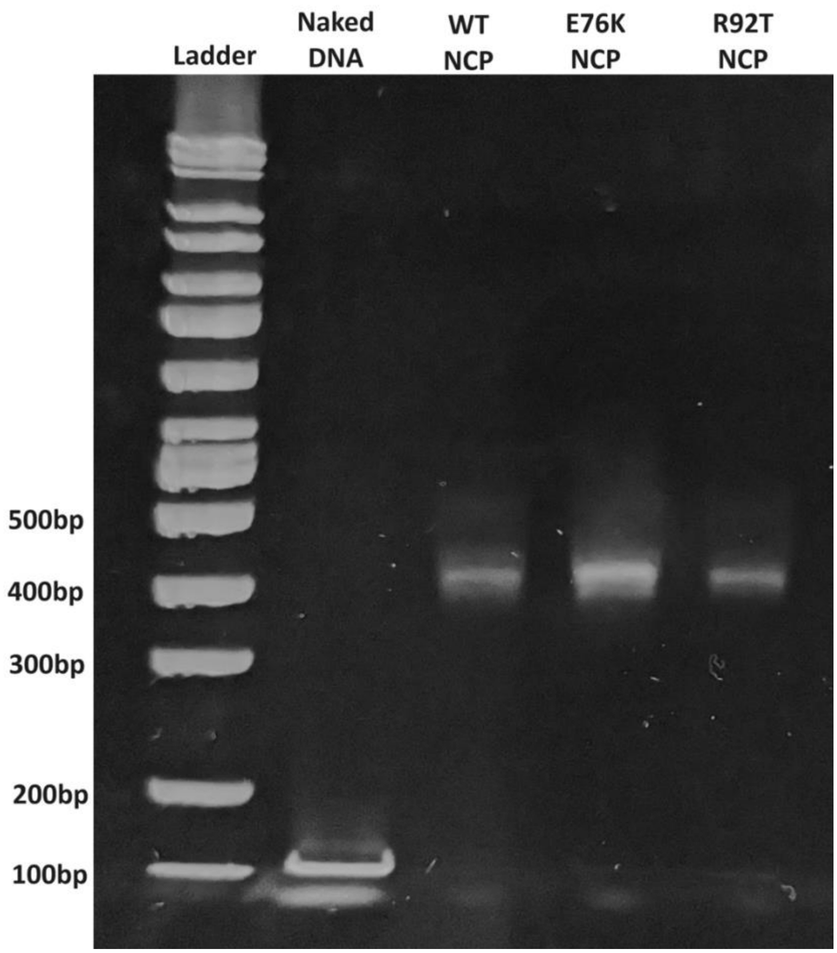


**Suppl. Figure 15:** 5% Acrylamide EMSA stained with ethidium bromide.

1. Lehmann, K., R. Zhang, N. Schwarz, A. Gansen, N. Mücke, J. Langowski, and K. Toth. 2017. Effects of charge-modifying mutations in histone H2A α3-domain on nucleosome stability

assessed by single-pair FRET and MD simulations. *Scientific Reports*. 7(1), doi: 10.1038/s41598-017-13416-x, https://dx.doi.org/10.1038/s41598-017-13416-x.
